# A role for a micron-scale supramolecular myosin array in adherens junction cytoskeletal assembly

**DOI:** 10.1101/2021.03.12.435158

**Authors:** Hui-Chia Yu-Kemp, Rachel A. Szymanski, Nicole C. Gadda, Madeline L. Lillich, Mark Peifer

## Abstract

Epithelial cells assemble specialized actomyosin structures at E-Cadherin-based cell-cell junctions, and the force exerted drives cell shape change during morphogenesis. The mechanisms used to build this supramolecular actomyosin structure remain unclear. We used ZO-knockdown MDCK cells, which assemble a robust, polarized and highly organized actomyosin cytoskeleton at the zonula adherens, and combined genetic and pharmacological approaches with super-resolution microscopy to define molecular machines required. To our surprise, inhibiting individual actin assembly pathways (Arp2/3, formins or Ena/VASP) did not prevent or delay assembly of this polarized actomyosin structure. Instead, as junctions matured, micrometer-scale supramolecular myosin arrays assembled, with aligned stacks of myosin filaments adjacent to the apical membrane, while associated actin filaments remained disorganized. This suggested these myosin arrays might bundle actin at mature junctions. Consistent with this, inhibiting ROCK or myosin ATPase disrupted myosin localization/organization, and prevented actin bundling and polarization. These results suggest a novel mechanism by which myosin self-assembly helps drive actin organization to facilitate cell shape change.

**Summary:** We explored mechanisms epithelial cells use to assemble supramolecular actomyosin structures at E-Cadherin-based cell-cell junctions. Our data suggest individual actin assembly pathways are not essential. Instead, microscopy and pharmacological inhibition suggest micrometer-scale supramolecular myosin arrays help bundle actin at mature junctions.

## Introduction

E-Cadherin (Ecad)-based adherens junctions (AJs) play a pivotal role in maintaining epithelial tissue homeostasis by mediating cell-cell adhesion and anchoring the actomyosin cytoskeleton (Lecuit and Yap, 2015). During morphogenesis, AJs ensure tissue integrity when cells change shape, divide, and move, events involving force exerted on AJs, which must be remodeled to accommodate tissue-wide mechanical forces. The relationship between AJs and the actomyosin cytoskeleton is one of reciprocal reinforcement (Michael and Yap, 2013). Apical AJs polarize actin, and connections to actin stabilize AJs at the plasma membrane. This focused attention on the complex supramolecular actomyosin structures that assemble at adherens junctions.

One key issue in the field is to define mechanisms cells use to assemble and polarize this complex structure. Significant progress has been made in examining polarization of AJs and the junctional cytoskeleton. *Drosophila* embryogenesis provides an important model, as 6000 cells simultaneously form and polarize their AJs during cellularization (Schmidt and Grosshans, 2018). Gastrulation then begins within minutes, and requires intricate cross-talk between AJs and the cytoskeleton, allowing dramatic cell shape changes while maintaining tissue integrity. However, the complexity of the *in vivo* system provides major challenges. For example, myosin, its activator Rho (e.g. (Crawford et al., 1998; Royou et al., 2004; Xue and Sokac, 2016), Arp2/3, and the formin Diaphanous (Afshar et al., 2000; Stevenson et al., 2002; Zallen et al., 2002) are required for cellularization. Cultured mammalian cells provide a simpler system in which to examine how the actomyosin cytoskeleton is assembled at AJs, with pharmacological tools allowing disruption of protein function in a temporally-controlled way. Often the junctional cytoskeleton is polarized, with Ecad and actin enriched at the apical end of lateral cell borders, in a structure known as the zonula adherens (ZA).

Scientists have taken several approaches to explore molecular mechanisms involved in ZA assembly and maintenance. One approach was to explore roles of actin and its regulators. Actin filament assembly involves nucleation, elongation, and bundling. The Arp2/3 complex nucleates new filaments from the sides of existing ones, creating branched networks, and formin family members nucleating new unbranched filaments (Buracco et al., 2019; Pollard, 2016). Formins and Ena/VASP proteins promote filament elongation. Blocking actin polymerization using cytochalasin or latrunculin can block de novo AJ assembly (Ivanov et al., 2005), but mature AJs are more resistant (Ivanov et al., 2004) (Tang and Brieher, 2012). The Arp2/3 complex coIPs with Ecad (Kovacs et al., 2002), and is enriched at AJs in many epithelial cell types (Yamada and Nelson, 2007). RNAi knockdown (Verma et al., 2012) or use of pharmacological inhibitors (Kovacs et al., 2011; Tang and Brieher, 2012) revealed a role for Arp2/3 in maintaining actin levels at AJs, but inhibiting Arp2/3 reduced but did not eliminate junctional actin. Multiple Arp2/3 activators, including cortactin (Han et al., 2014; Helwani et al., 2004), N-WASP, WAVE, WIRE, (Verma et al., 2004) (Kovacs et al., 2011; Verma et al., 2012) also are enriched at AJs and interact with Ecad. However, once again loss-of-function reduced but did not eliminate junctional actin in established monolayers, suggesting Arp2/3 acts in parallel with other mechanisms to assemble junctional actin. Scientists also explored formins (Carramusa et al., 2007; Kobielak et al., 2004; Nishimura et al., 2016; Rao and Zaidel-Bar, 2016; Sahai and Marshall, 2002). Individual knockdown of DAAM1 or Dia1 reduced but did not eliminate junctional actin. Finally, scientists explored Ena/VASP proteins. While Ena/VASP proteins are enriched at AJs (e.g., (Oldenburg et al., 2015; Vasioukhin et al., 2000) knockdown or sequestration away from cell junctions reduced but did not eliminate junctional actin (Scott et al., 2006; Yu-Kemp et al., 2017). Taken together, these data suggest multiple parallel mechanisms drive assembly and maintenance of the junctional actin cytoskeleton.

Scientists have also explored roles of non-muscle myosin II (hereafter referred to as myosin). Junctional contractility at the ZA is driven by myosin and regulated by complex feedback loops between actin, myosin, and Ecad. Myosin monomers assemble into bipolar filaments and their ATPase-powered motor activity produces contractile force on the AJ-associated actin cytoskeleton (Agarwal and Zaidel-Bar, 2019; Vicente-Manzanares et al., 2009). Myosin filament assembly requires phosphorylation via Rho kinase (ROCK) or myosin light chain kinase (MLCK). Myosin also crosslinks actin, which, coupled with its motor activity, allows myosin to organize actin. Inhibiting myosin motor activity or myosin activation reduced, but did not eliminate, actin assembly at AJs (Leerberg et al., 2014; Sahai and Marshall, 2002; Shewan et al., 2005). Myosin-generated contractility can stimulate actin assembly (Leerberg et al., 2014), and myosin contractility can stabilize AJs by pulling on alpha-catenin and shifting it to an open state, strengthening AJ:actin connections (Ozawa, 2018)

Advanced imaging revealed new insights into supramolecular myosin organization. Two prescient papers from the 1990s explored myosin organization in the lamella of migrating fibroblasts. Myosin filaments can stack on one another, in arrays involving dozens of filaments (Svitkina et al., 1997; Verkhovsky et al., 1995). Structured illumination microscopy (SIM) allowed direct observation of myosin stacks and associated actin filaments in living cells (Fig 1A; (Beach et al., 2017; Fenix et al., 2016; Hu et al., 2017). Individual myosin filaments organized into stacks, with stacks oriented perpendicular to peripheral stress fibers, thus linking actin filaments (Fig 1A), and aligned myosin stacks formed two-dimensional arrays tightly apposed to the plasma membrane. Inhibiting myosin activation or myosin motor activity reduced stack assembly. Hu and Bershadsky made interesting prediction: “Since the interaction of myosin filaments associated with different actin bundles creates forces attracting these bundles towards each other, the organization of myosin filaments into stacks is a plausible mechanism for the formation of the densely packed arrays of parallel actin bundles often observed in polarized fibroblast-type cells” (Hu et al., 2017). Advances in imaging also provided insights into actin and myosin organization at the ZA, revealing two zones of actin at the ZA of endothelial cell bicellular borders (Efimova and Svitkina, 2018): a central zone of branched actin immediately adjacent to Ecad, co-localizing with Arp2/3, and more lateral bundled actin filaments decorated by myosin, forming an actin belt along each bicellular border. Superresolution microscopy is consistent with similar organization of actin and myosin at bicellular junctions in other epithelial cells (Choi et al., 2016; Heuze et al., 2019; Kovacs et al., 2011). At tricellular junctions, bundled bicellular border actin filaments may anchor end on in cadherin-catenin complexes (Choi et al., 2016; Efimova and Svitkina, 2018).

**Fig 1.**
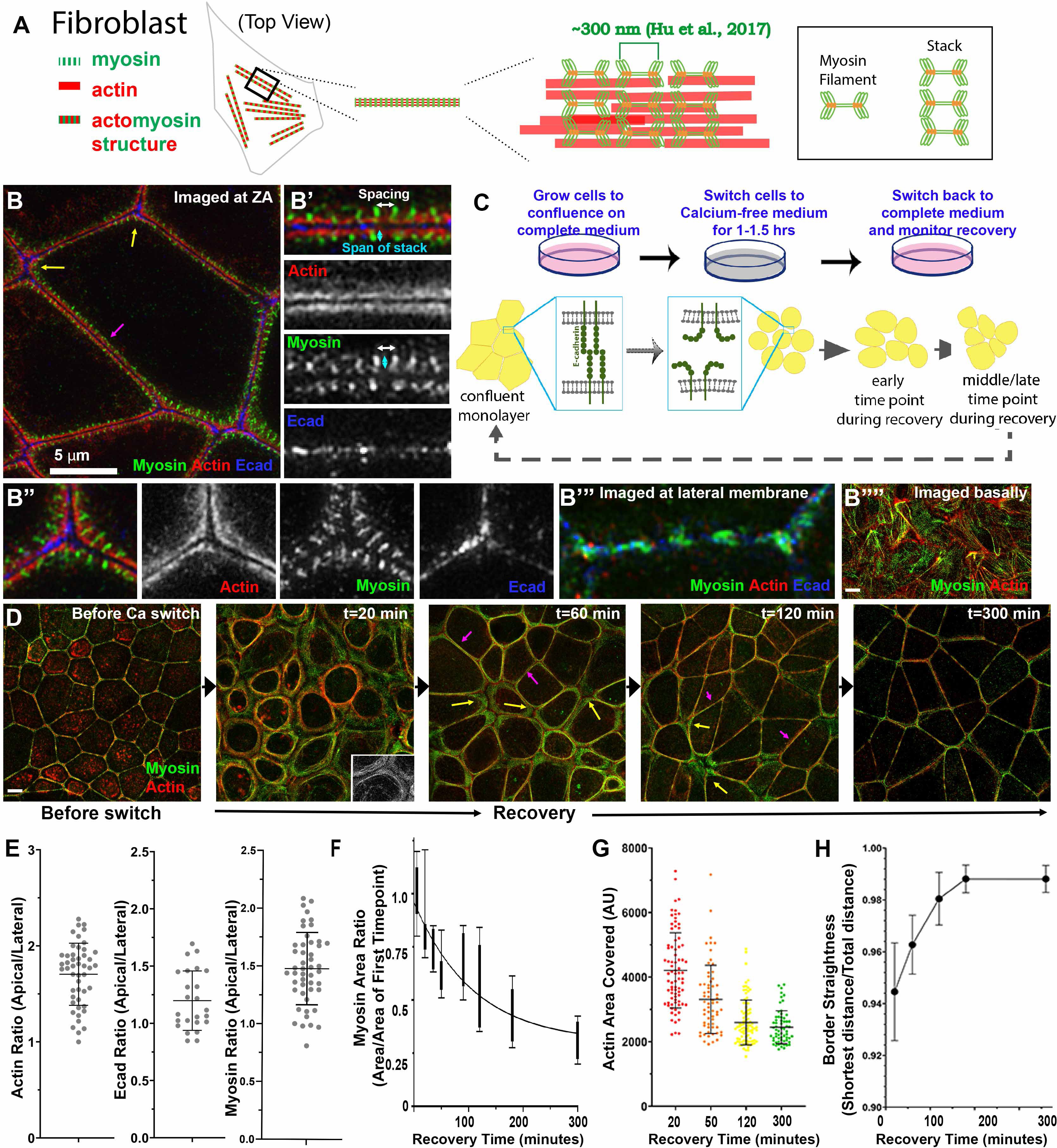
ZO KD MDCK cells as a model to study formation of ZA actomyosin structures. (A) Schematic of myosin filament stacks observed in fibroblasts. (B) ZO KD MDCK cells. ZA at bicellular borders (B’) and tricellular junctions (BA’’). Actin is bundled and myosin is organized into a sarcomeric pattern. (B’’’) Lateral membrane. (B’’’) basal stress fibers. (C) Schematic diagram of calcium switch experiments. (D) Representative images of actin and myosin at the apical surface as junctions recover. (E) Ratio of apical to lateral signals of actin, E-Cad or myosin at the end of recovery. (F-H) Quantification. Changes of area covered by myosin (E), ZA actin bundling (F), and border curvature (H) during junction maturation. for all the figures, unless indicated. For all figures, unless otherwise indicated: Top-view images are apical maximum intensity projections (MIP); arrows: (magenta) bicellular borders, (yellow) tri/multicellular junctions; scale bars=5 µm

To define mechanisms by which cells assemble and position the contractile cytoskeleton at the ZA as AJs assemble, we combined a model epithelial cell line that assembles a robust actomyosin array at the ZA with superresolution imaging to determine the ZA’s organization and assembly pathway (Choi et al., 2016; Fanning et al., 2012). ZO-knockdown MDCK cells assemble an “Albert’s textbook” ZA, much like the ZA sarcomeric array of cochlear hair cells (Ebrahim et al., 2013). We used the calcium switch assay to allow us to visualize reassembly of the supramolecular actomyosin array at the ZA via super-resolution microscopy, and combined this with genetic and pharmacological approaches to define the molecular machines and mechanisms involved.

## Results

### A model system to study actomyosin assembly at the zonula adherens

ZO-knockdown MDCK cells (Choi et al., 2016; Fanning et al., 2012) assemble a textbook-like ZA with a highly organized actomyosin array positioned at the apical end of lateral cell borders (Fig 1B). SIM super-resolution imaging revealed tightly bundled actin cables along bicellular junctions (Fig 1B, magenta arrow; B’), decorated by a sarcomeric array of myosin (Fig 1B white arrow=sarcomeric spacing) and underlain by puncta of E-cadherin (Ecad; (Choi et al., 2016);Fig 1B’). There are elevated levels of Ecad and increased spacing between actomyosin arrays at tricellular junctions (Fig 1B, yellow arrows, 1B”), where molecular tension is exerted on cadherin-catenin complexes (Choi et al., 2016). Ecad is enriched at the apical ZA (Suppl Fig 1A, red arrows; quantified in Fig 1E) and also present at lower levels all along lateral cell borders (Suppl Fig 1A, green arrows), where it associated with a distinct actin population that is less bundled and had little associated myosin (Fig 1B”’). Actin and myosin are enriched at the apical ZA (Fig 1E), and are also found in basal stress fibers (Fig 1B””).

To study AJ re-assembly, we used the well-characterized calcium switch assay (Gumbiner and Simons, 1986). Before perturbation cells were columnar with a slightly domed apical microvillar surface. We removed calcium for 1.5 hours, disrupting cell-cell adhesion, and cells went from columnar to rounded (Fig 1D, 0 min to 20 min; Suppl Fig 1F). Ecad was endocytosed, accumulating in an apparent vesicular-compartment (Suppl Fig 1A, cyan arrows) and cells unzipped along their lateral borders (magnesium was not chelated, so cell-ECM adhesion was not disrupted). When calcium was added back, Ecad returned to the bases of the cells where contact was preserved (Suppl Fig 1, red arrows)—at this stage cells were highly rounded (Suppl Fig 1F, M). As AJs reassembled over several hours (Fig 1D), cells zipped together, with Ecad going from punctate at cell junctions to more continuous at re-established junctions (Suppl. Fig 1B vs C,D, red arrows) (Suppl Fig 1C,D, red arrows). Over the next 2-3 hours, borders slowly straightened; Bicellular borders straightened first (Fig 1D, magenta arrows), with tricellular and short multi-cellular junctions the last sites of reassembly (Fig 1D, Suppl Fig 1C, yellow arrows), until columnar architecture was restored (Suppl Fig 1F-Q), with Ecad enriched at the ZA (Suppl Fig 1E, red vs green arrows). We adopted three methods to quantify the rate of junctional actomyosin reassembly. First we quantified the area occupied by myosin over the entire field. As AJs re-assembled, myosin arrays at junctions narrowed, and thus the area occupied by myosin decreased until it reached that seen before calcium withdrawal (Fig 1F). Similarly, we quantified increased actin bundling into the tight array seen before perturbation, by selecting apical regions of bicellular junctions, binarizing images, and calculating the area occupied by actin (Fig 1G). Finally, we quantified junctional straightening as AJs reassembled (Fig 1H). These data were the baseline for our subsequent perturbations.

### Junctional reassembly involves generation of very large-scale myosin arrays

We next examined AJ and cytoskeletal proteins at high resolution as AJs re-assembled, using Zeiss Airyscan or SIM imaging. Myosin and actin localization proved quite surprising. Before perturbation, myosin localized to the tight ZA sarcomeric array (Fig 1B-B”), and to basal stress fibers (Fig 1B””). As cells rounded up after calcium removal, myosin initially associated with actin “arcs” aligned along reforming lateral cell borders (Fig 2A-A’”; closeup in 2B). However, strikingly, within 20 min of recovery, extensive stacks of myosin filaments assembled around the cell periphery (Fig 1D, 20 min; Fig 2C), similar to but more extensive than the myosin stacks observed in migrating fibroblasts (Beach et al., 2017; Fenix et al., 2016; Hu et al., 2017; Verkhovsky et al., 1995). These myosin stacks rapidly returned to their mature form at many bicellular borders (Fig 1D 60 min; Fig 2C, magenta arrows) but remained extensive at a subset of tricellular and multicellular borders as recovery proceeded (Fig 1D 60 min; Fig 2C, yellow arrows).

**Fig 2.**
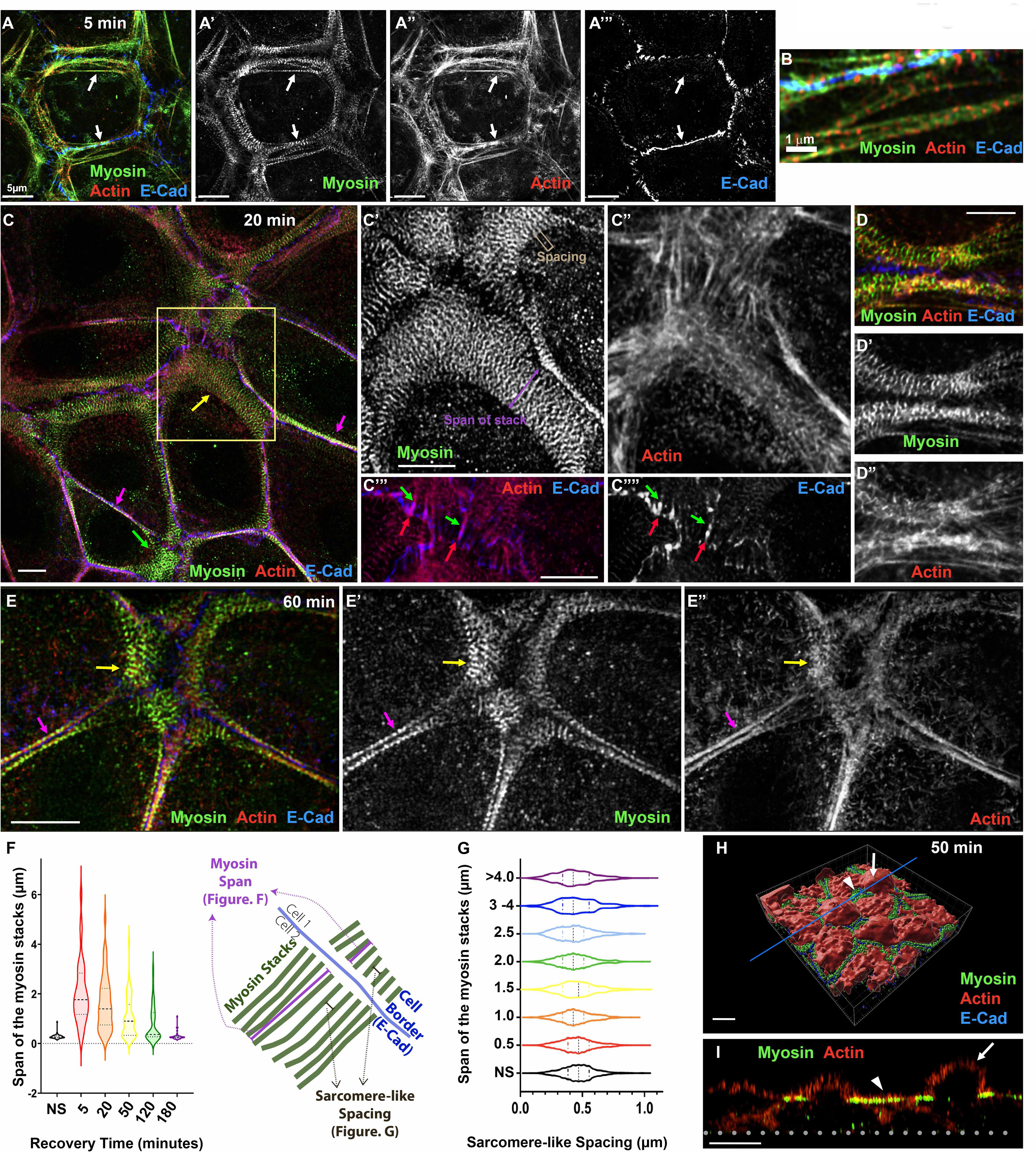
Extensive stacks of myosin precede the formation of bundled actin as E-cad based junctions assemble. (A-E) Images collected at different time points of calcium recovery reveal that an extensive myosin array forms during mid-recovery and localizes with a less well organized actin network. (F-G) Quantification. Despite changes in the span of the myosin stacks at different time points (F), spacing between the myosin stacks remains similar (G). (H) 3-D. surface rendering of a cell in mid-recovery. (I) Cross-section view at the line indicated in H. The extensive myosin stacks localize to a restricted Z plane underlying the apical plasma membrane (arrowheads), between the domed microvillar caps (arrows).

The micrometer-scale supramolecular myosin arrays assembled during recovery were quite striking. In unperturbed cells, the sarcomere-like myosin arrays decorating the bundled actin filaments spanned ∼200-300 nm in the direction perpendicular to the membrane (Fig 1B’, blue arrow; Fig 2F NS). In contrast, during mid-assembly stacks could exceed many times that span, extending to 4-6 µm (Fig 2C,C’ purple arrow; Fig 2F). In unperturbed cells, myosin at the ZA had a spacing between the short myosin filament stacks of ∼400nm (Fig 1B’, white arrow; (Choi et al., 2016); the antibody detects myosin heavy chain’s tail), similar to spacing in stress fibers. Strikingly, spacing between stacks remained similar throughout assembly (Fig 2C’, bracket), regardless of the span of the stacks (Fig 2G) or age of the junction (Suppl Fig 1L). Actin underlying these myosin stacks was less organized than myosin, often forming a meshwork (Fig 2C”). There also were actin structures associated with Ecad at reforming junctions, with short robust actin filaments (Fig 2C’”, C””, red arrows) terminating in interdigitating cadherin-zippers (Fig 2C’”, C”” green arrows), like those seen in keratinocytes or endothelial cells (e.g. (Huveneers et al., 2012; Vasioukhin et al., 2000). At times, remnant actin “arcs” parallel to the lateral membrane were seen underlying the myosin stacks (Fig 2D-D”), similar to those observed earlier in recovery (Fig 2A-A”). Myosin arrays at bicellular and some tricellular junctions rapidly reduced their span, but at a subset of tricellular and multicellular junctions extensive myosin stacks remained and actin in these regions remained less well organized (Fig 2E). These were the last places to resume the bundled actin architecture seen before the calcium switch (Fig 1D 120 min, yellow arrows).

We next examined myosin stack positioning along the z-axis. Strikingly, myosin stacks occupied an ∼0.5iilm region in the Z plane, close to and underlying a flattened region of the plasma membrane “apical” to lateral borders (Fig 2H-I’, arrowheads; Suppl. Fig 1M) and separate from the domed center (Fig 2H-I, arrows). The domed membrane flattened as AJs matured, but residual myosin stacks continued to occupy flattened regions underlying the apical membrane throughout the process (Suppl. Fig 1K, M-Q). Together, these data reveal that very extensive myosin stacks form during ZA reassembly, often before there is any apparent organization of underlying actin, and these stacks narrow as actin is bundled into tight polarized sarcomeric structures, reforming the ZA.

### Neither Arp2/3 nor formin activity are essential to assemble specialized actomyosin structures at the ZA, although they do enhance cortical actin

We next used functional assays to define mechanisms driving assembly of specialized ZA actomyosin structures. We tested two broad mechanistic hypotheses: 1) ZA actomyosin assembly is driven by actin filament nucleation and elongation, or 2) ZA actomyosin assembly is driven by myosin’s motor activity, gathering and bundling pre-existing actin filaments. We initially favored a role for the Arp2/3 complex, as it can coIP with Ecad, localizes to AJs, can stimulate actin assembly at Ecad-based contacts, and along with its regulators helps stimulate junctional actin assembly in confluent monolayers (Efimova and Svitkina, 2018; Kovacs et al., 2011; Tang and Brieher, 2012; Verma et al., 2012). We first examined Arp2/3 localization during junction recovery. As in other cell types, the Arp2/3 complex was enriched at the ZA in unperturbed confluent cells (Fig 3A arrow). However, enrichment was weak until relatively late in junctional reassembly (Fig 3B-E).

**Fig 3.**
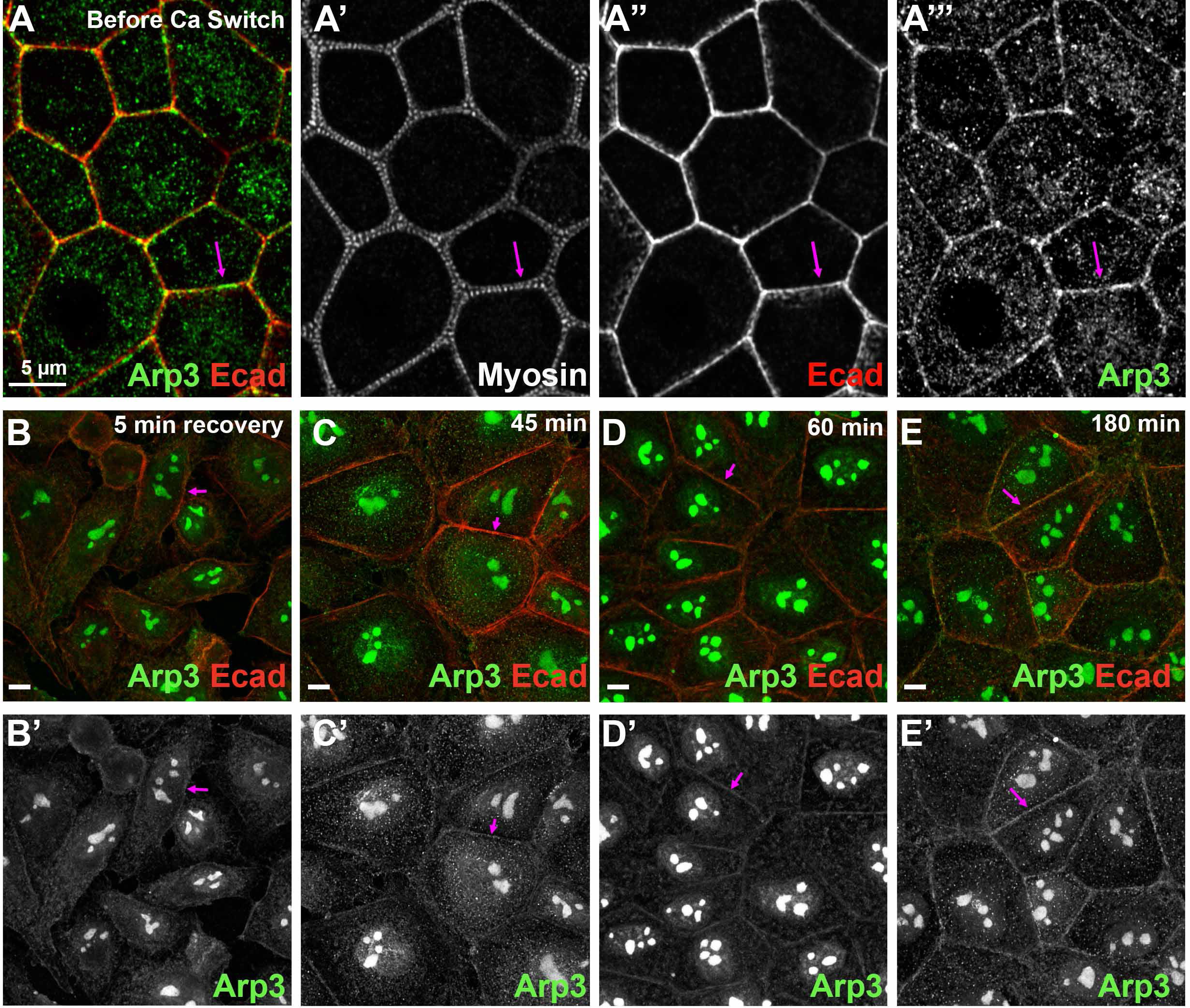
The Arp2/3 complex localizes to the ZA in confluent ZO KD MOCK cells but only returns there slowly after calcium switch. (A) Arp3 at AJs in a confluent, mature monolayer. (B-E) Arp3 is only weakly localized to junctions during early stages of recovery.

To assess Arp2/3’s role, we used the well characterized inhibitor CK666 (Hetrick et al., 2013; Nolen et al., 2009), after verifying its activity by assessing its ability to block cell spreading on the substrate (Suppl Fig 2). We carried out the calcium switch in the presence or absence of CK666. To our surprise, assembly of the tight ZA actomyosin array was qualitatively unaltered by the inhibitor (Fig 4A-C vs D-F). Cells rounded up at early time points (Fig 4A vs. D), and then expansive myosin stacks formed as junctions were re-established (Fig 4B vs E, yellow arrows). These stacks slowly reduced in span, first at bicellular junctions (Fig 4B vs E, magenta arrows) and finally at tricellular junctions, until the ZA sarcomeric actomyosin array was re-established (Fig 4C vs F). Super-resolution imaging confirmed that in places with extensive myosin stacks actin was present as a disordered array of filaments (Fig 4J, J inset arrows), as in controls. In CK666 treated cells, tightly bundled actin and narrowed stacks of sarcomeric myosin were restored at the ZA at both bicellular and tricellular junctions (Fig 4K, magenta and yellow arrows). To assess whether junctional re-establishment was delayed after Arp2/3 inhibition, we used reduction in span of myosin stacks to quantitatively assess these events (as in Fig 1F). While there was experiment to experiment variability, there was no substantial delay myosin stack narrowing after Arp2/3 inhibition (Fig 4L). Intriguingly, Arp2/3 inhibition reduced total apical actin levels and modestly reduced lateral actin (Fig 4M), but apical actin enrichment at the ZA was unaffected (Fig 4N). We assessed ZA actin bundling (as in Fig 1G). At the completion of ZA reassembly, actin was equally bundled after Arp2/3 inhibition (Fig 4O). Finally, border straightening occurring as the ZA actomyosin array assembled was also unaltered (Fig 4P). Together, these data suggest Arp2/3 activity is not essential for timely and accurate assembly of specialized ZA actomyosin structures, even though it promotes overall cortical Actin levels in this cell type.

**Fig 4.**
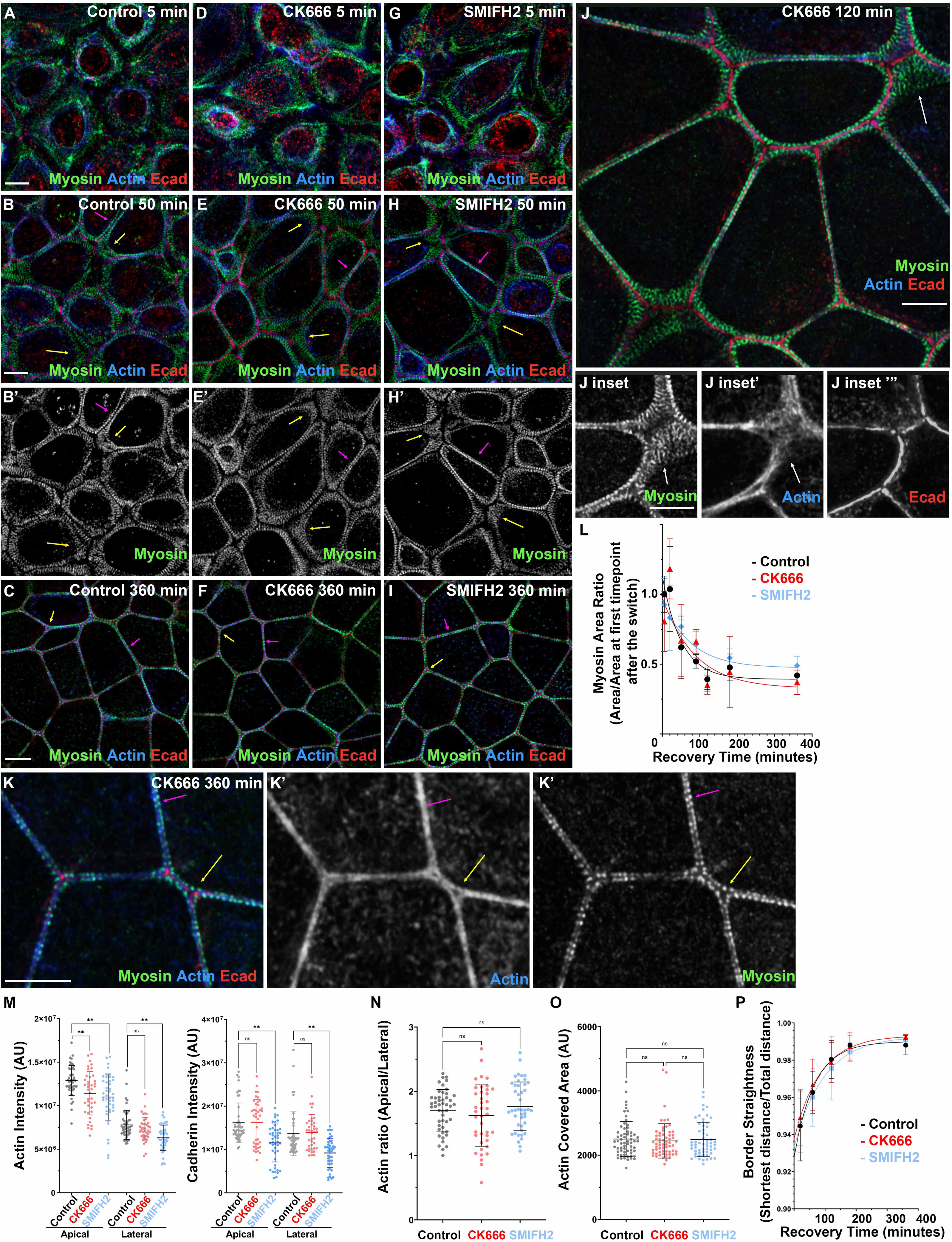
Neither Arp2/3 nor formin activity are essential to assemble specialized actomyosin structures at the ZA, although they do enhance cortical actin. (A-K) Recovery from calcium switch in control, CK666 (Arp2/3inhibitor)-treated, or SMIFH2 (formin inhibitor)-treated cells. (L-P) Quantification. Narrowing of the myosin stacks (L), actin bundling (O), and cell border straightening (P) remain similar among all three conditions. However, inhibition of actin nucleators reduce apical actin levels (M, left), but parallel changes in lateral actin mean that apical polarization of actin is not altered (N). **: p<0.01; ns=non-significant.

Formins have also been implicated in AJ assembly/maintenance (Carramusa et al., 2007; Kobielak et al., 2004; Nishimura et al., 2016; Rao and Zaidel-Bar, 2016; Sahai and Marshall, 2002). Mammals have many formins, rendering knockdown approaches challenging. We thus used a formin FH2 domain inhibitor, SMIFH2 (Rizvi et al., 2009), at a concentration like that used by others (50µM), in the calcium switch to determine its effect on actomyosin architecture. As with Arp2/3 inhibition, we observed no substantial qualitative (Fig 4A-C vs G-I) or quantitative differences (Fig 4L-P) in assembling specialized actomyosin structures at the ZA. Formin inhibition did lower levels of both apical and lateral actin (Fig 4M), as others observed (e.g. (Rao and Zaidel-Bar, 2016), but did not alter its apical polarization at the ZA (Fig 4N). We also noted what appeared to be cell toxicity at later time points, in which cells were lost in small regions of the field (Suppl. Fig 3)—this may be due to off-target effects. It is important to note that the specificity of this inhibitor has come into question, with evidence that it can also inhibit myosin 2 (Nishimura et al., 2021). However, since SMIFH2 treatment did not alter ZA actomyosin assembly, while myosin inhibition did (see below), we do not think those particular off-target effects alter our conclusions. Thus, with those caveats, there was no requirement for formin activity in assembling specialized ZA actomyosin structures.

### Sequestering Ena/VASP proteins does not block or slow assembly of specialized actomyosin structures at the ZA

Ena/VASP proteins bind to the growing ends of actin filaments to antagonize capping and stimulate monomer addition (Bear and Gertler, 2009). In some cell types Ena/VASP proteins are implicated in AJ assembly/maintenance (Scott et al., 2006; Yu-Kemp et al., 2017). We first examined VASP localization in our cell line. In confluent, unperturbed cells, VASP was enriched at the ZA, with special enrichment at tricellular junctions (Fig 5A, arrows), thus matching Ena localization in *Drosophila* tissues (Gates et al., 2007; Rauskolb et al., 2019). A different fixation technique increased junctional signal, and, because AJs separated a bit, revealed VASP localizes slightly membrane proximal to the actomyosin array (Fig 5B, arrows). We then explored how VASP localization changes during junctional reassembly. Right after calcium switch, VASP primarily localized basally to the ends of stress fibers (Fig 5C, arrows). Midway through recovery VASP localized between the expanded myosin stacks with some enrichment at tricellular/multicellular junctions (Fig 5D,E). This was consistent with a potential role in actomyosin assembly at the ZA.

**Fig 5.**
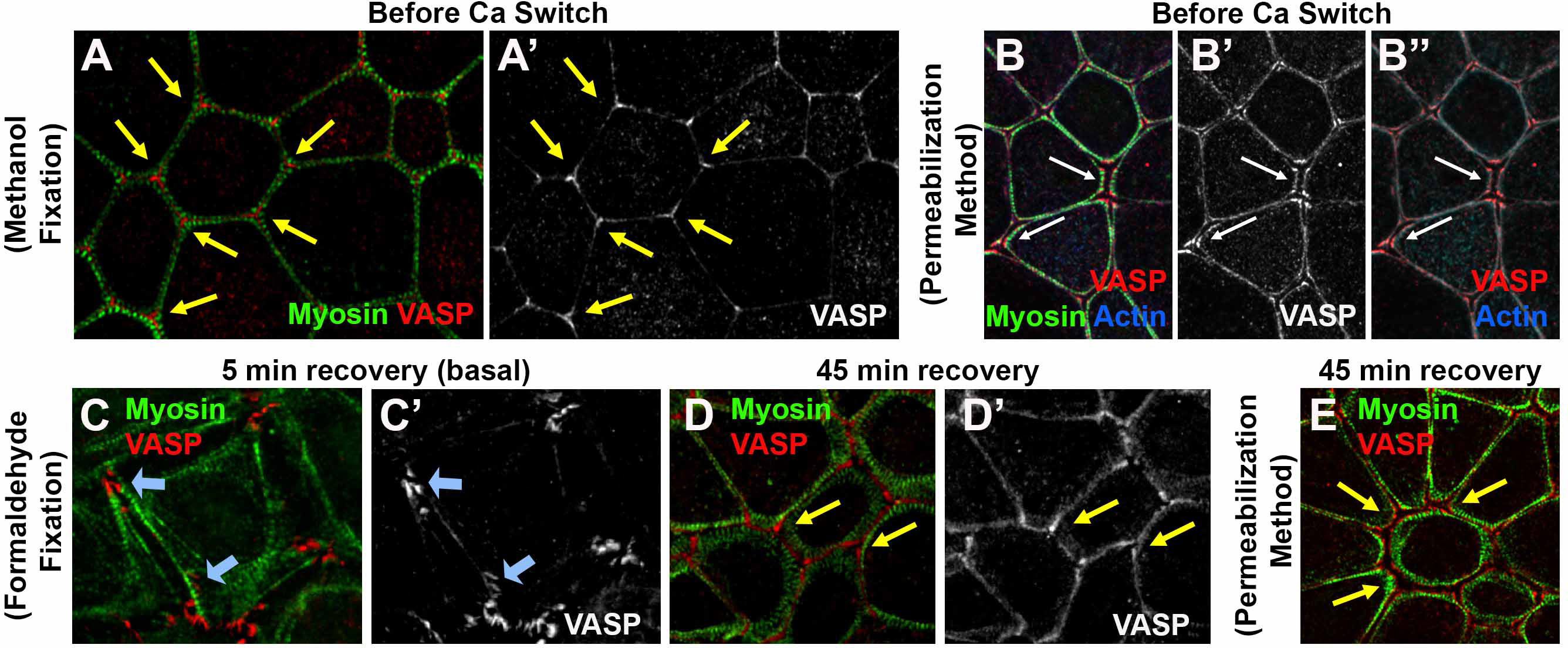
VASP localizes to cell borders with enrichment at tricellular junctions. (A,B) In confluent monolayers VASP localizes to bicellular borders and is enriched in tricellular junctions. (C-D). At early stages of recovery, VASP also localizes to focal adhesions (C). As E-cad junctions start to form, VASP is detected at cell borders and tricellular junctions.

The presence of three Ena/VASP family members complicates knockdown, but a clever approach allows sequestration of all three at mitochondria, thus inactivating them. FP4mito fuses a sequence that inserts in the mitochondrial outer membrane with sequences encoding high affinity Ena/VASP binding sites, while in the control construct, AP4, the Ena/VASP sites are mutated (Bear et al., 2000). We confirmed that FP4mito sequestered VASP in our cell line (Suppl Fig 2F), and used it to test whether Ena/VASP proteins play an important role in assembling specialized ZA actomyosin structures.

To test whether Ena/VASP proteins play such a role, we transfected cells with FP4mito, or AP4mito as a control, and subjected them to calcium switch (Fig 6). Transfection efficiency was not 100% so we focused on regions where several FP4mito or AP4mito-expressing cells were adjacent (FP4mito or AP4mito-expressing cells are indicated by asterisks). Surprisingly, sequestering Ena/VASP proteins did not affect ZA actomyosin array assembly, as this proceeded in a similar way in FP4mito and control AP4mito cells. Cell rounded up after calcium withdrawal, with myosin and actin predominantly in basal stress fibers (Fig 6A vs B, arrows). Midway through recovery, broad myosin stacks were observed in both cell populations (Fig 6C vs D, arrows insets), with narrowing occurring first at a subset of bicellular junctions. The myosin stacks were qualitatively similar to those observed in untransfected cells. Finally, expressing FP4mito did not alter the final ZA sarcomeric myosin array decorating tightly bundled actin filaments (Fig 6E vs F, arrows insets). Quantitative analysis of myosin stack narrowing revealed no delay after FP4 transfection relative to AP4 (Fig 6G). In contrast to inhibiting Arp2/3 or formins, sequestering Ena/VASP did not reduce overall cortical actin levels (Fig 6H). Actin apical polarization was unaffected (Fig 6I), as was actin bundling at the ZA (Fig 6J). Finally, Ena/VASP sequestration did not slow junction straightening (Fig 6K). Thus Ena/VASP proteins do not appear to be essential for assembling the highly ordered ZA actomyosin array.

**Fig 6.**
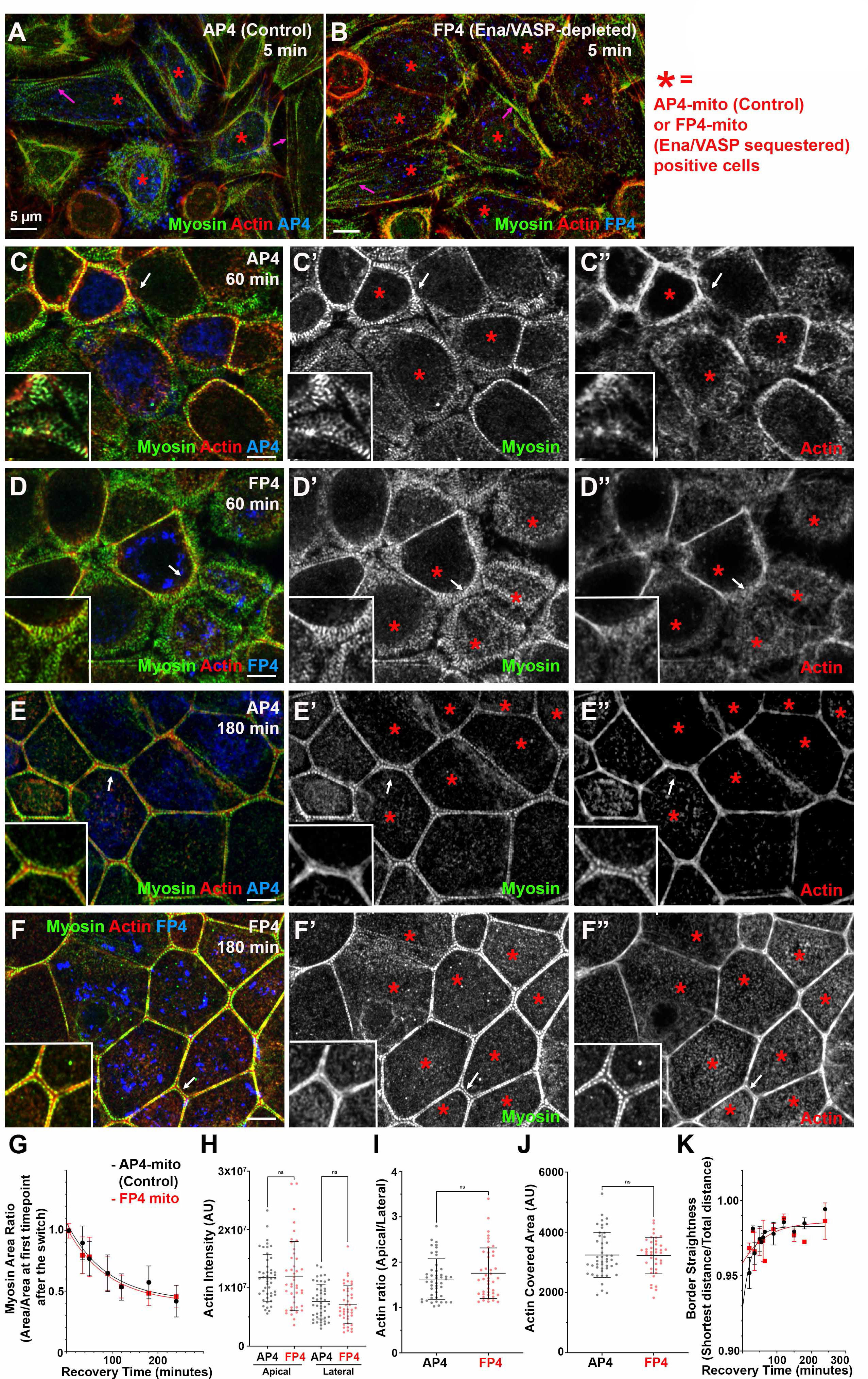
Ena/VASP proteins are not essential for assembly of specialized actomyosin structures at the ZA. (A-F) Representative images, different timepoints during recovery. Control (AP4) or VASP-sequestered (FP4). (G-K) Quantification. There was no difference between control and VASP-sequestered cells in narrowing of myosin arrays (G), actin intensity (H), apical actin polarization (I) actin bundling (J), or junction straightening (K). White arrows: locations ofthe zoom-in insets.

### Proper assembly of tightly bundled actin at the ZA requires ROCK activity but not MLCK

The data above suggest individual actin polymerization machines are not essential for assembling the highly organized ZA actomyosin structures. We next tested our second hypothesis: ZA actomyosin assembly is driven by myosin motor activity, gathering and bundling pre-existing actin filaments. The expansive myosin stacks assembled as junctions reformed and actin became bundled were consistent with that hypothesis. We first examined kinases that phosphorylate and activate myosin: Rho kinase (ROCK) and Myosin Light Chain Kinase (MLCK). We used well characterized, specific inhibitors of each to explore roles of ROCK and MLCK

To test if MLCK activation of myosin is essential to drive ZA actomyosin assembly, we incubated cells with ML-7, which inhibits MLCK’s catalytic activity (Saitoh et al., 1987), during recovery after calcium switch. After ML-7-treatment, myosin still localized to cell borders. There was no noticeable delay in myosin stack narrowing or junctional actin bunding as junctions matured, compared to control (Suppl Fig 4A-F). Quantification confirmed that ML-7 treated cells had similar rate of myosin stack narrowing (Suppl Fig 4G) and ZA actin bundling (quantified in Suppl Fig 4I). Borders straightened at the same rate as in controls (quantified in Suppl Fig 4J). ML-7 treated cells had somewhat elevated actin levels laterally, thus reducing apical enrichment (quantified in Suppl Fig4K-L). We did less extensive experiments with a second MLCK inhibitor, peptide-18 (Lukas et al., 1999), and saw a similar lack of effect (Suppl Fig 4H; quantified in Suppl Fig 4M). These data suggest MLCK signaling is not essential for assembling actomyosin structures at the ZA.

We next tested whether ROCK has a role, using a well-characterized ROCK inhibitor Y-27632 (Ishizaki et al., 2000) during recovery. The result was quite different. ROCK inhibition reduced overall cortical myosin levels (Suppl Fig 4 N vs O, P vs Q) and disrupted the organized myosin stacks seen in controls—this was seen even at our earliest time points (Fig 7A vs B, arrows) and was reflected in the reduced area covered by myosin through most of the time course (Fig 8A) In controls, extensive myosin stacks appeared at tricellular and multicellular junctions by 20 min of recovery (Fig 7C, arrows; Fig 7C’” =magnification of the sarcomere-like array). In contrast, after ROCK inhibition, while some myosin accumulated overlapping cortical actin near junctions (Fig 7D, arrows), organized myosin stacks were absent. In controls myosin stacks at the ZA narrowed by 105 min, with a tight sarcomeric array and bundled actin at many bicellular borders (Fig 7E, magenta arrows), though a subset of tricellular junctions retained more extensive myosin stacks and less organized actin (Fig 7E, yellow arrows). By 180 min, controls returned to the highly organized bundled actin and sarcomeric myosin seen prior to perturbation (Fig 7G). In contrast, while actin accumulated at cell junctions in ROCK inhibitor treated cells, it did not form the tight bundled arrays seen in controls (Fig 7E vs F, G vs H, magenta arrows; quantified in Fig 8B), even at the latest timepoints (Fig 7H). Borders did not become straight, even at time points where recovery was complete in controls (Fig 7G vs H, arrows; quantified in Fig 8D). ROCK inhibition also reduced apical polarization of actin (Fig 7E’” vs F’”, apical-red arrows vs lateral-green arrows; Fig 7G’” vs H’”; actin quantified in Fig 8C), and also reduced overall levels of apical actin (Suppl Fig 4R). Interestingly, effects of ROCK inhibition were reversible, with tight sarcomeric myosin and bundled actin reforming after inhibitor washout (Fig 7I, magenta arrows; quantified in Fig 8A,B), and apical polarization re-established (Fig 7I inset, Fig 7I’”;Fig 8C). Thus myosin activation by ROCK is essential for assembly of sarcomeric myosin arrays and tightly bundled actin at the ZA.

**Fig 7.**
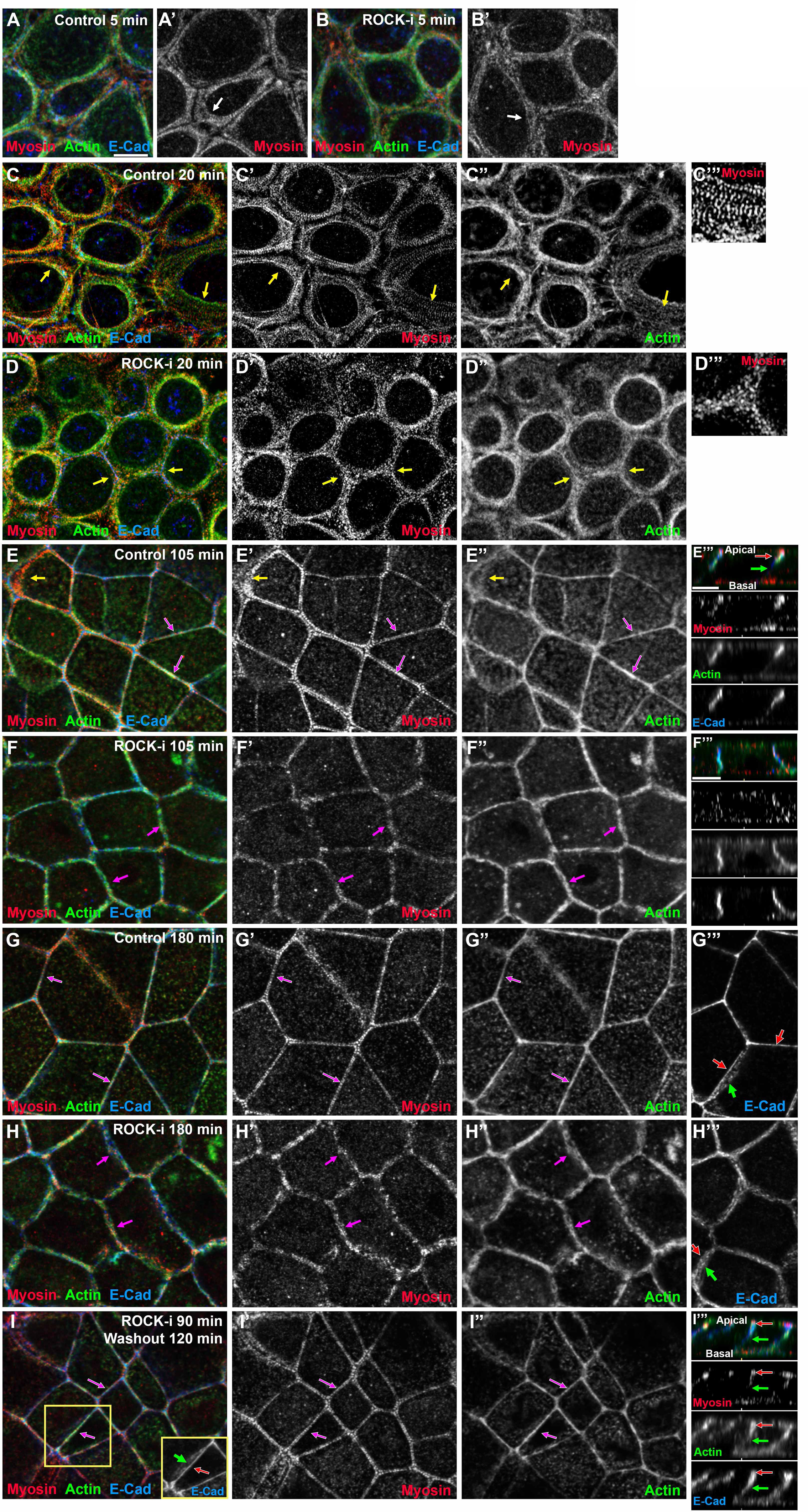
Inhibiting ROCK reduces myosin recruitment and organization at cell borders, leaving less bundled F-actin and less actin polarization at the ZA. (A-I) Representative images, different timepoints during recovery. Control vs. ROCK-inhibited (ROCK-i). ROCK inhibition reduces myosin at cell borders (Suppl Fig 4N-Q), but when its signal is intensified, it also becomes clear that myosin stacks are disrupted (C’ vs D’ arrows, C’” vs D’”) and at later timepoints sarcomeric organization ay the ZA is lost (E’ vs F’, G’ vs. H’). Actin is less bundled than control (E” vs F”, G” vs. H”). Apical polarization of Ecad, myosin and actin to the ZA is reduced after ROCK-i (E’” vs F’”, G’’’ vs. H’’’ red vs green arrows). (I-I’”) After inhibitor washout ZA actomyosin architecture is restored.

**Fig 8.**
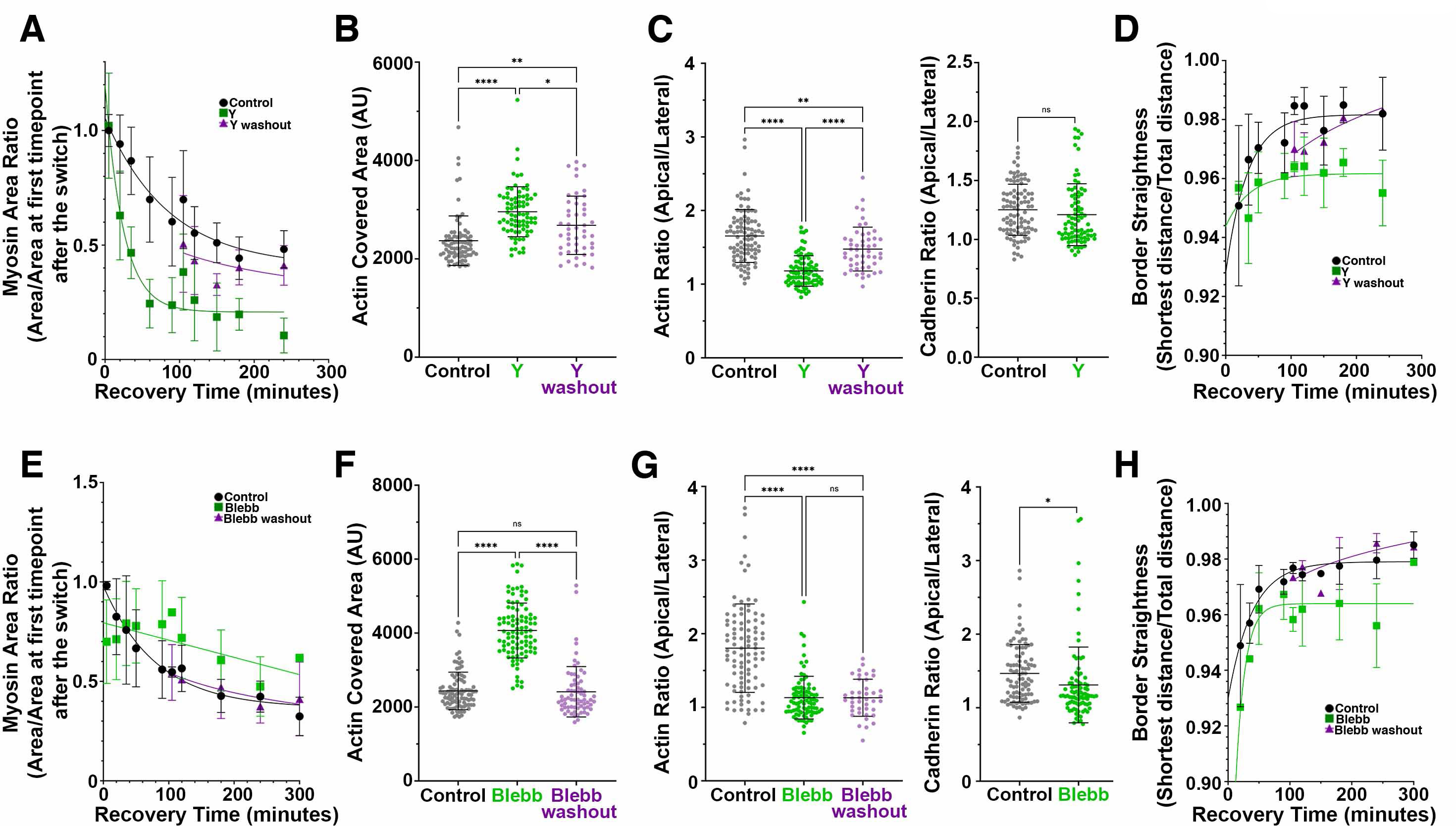
Myosin activation and motor activity are important for actin bundling and polarization at the ZA. Quantification. (A-D) ROCK inhibition. (E-H) Myosin-ATPase-inhibition. Inhibiting ROCK reduces junctional myosin (A) while blebbistatin reduces narrowing of myosin stacks (E). Both inhibitors reduce actin bundling at the ZA (B,F), apical actin polarization (C,G left), and border straightening (D, H).

### Proper assembly of tightly bundled actin at the ZA requires myosin motor activity

We next explored the role of myosin motor activity, which can play an important role in actin re-modeling. We used the inhibitor Blebbistatin, which inhibits myosin motor activity without abolishing its ability to bind actin (Kovacs et al., 2004). We repeated the calcium switch assay in the presence of this inhibitor. The results were interesting and surprising.

As expected, myosin was disrupted even at the earliest time points (Fig 9A vs B). Assembly of the extended myosin stacks seen in the controls (Fig 9C’, yellow arrows) was prevented by blebbistatin. Instead, myosin accumulated in a punctate pattern overlapping cortical actin (Fig, 9D’, D” yellow arrows), with some enrichment near Ecad-based cell junctions (Fig 9D, D’, magenta arrows). Cortical myosin remained disorganized even after extended recovery in the presence of blebbistatin (Fig 9F’, H’ red arrows), by which point control cells had resumed tight sarcomeric myosin arrays and bundled actin filaments and both bicellular and tricellular junctions (Fig 9G-G”, magenta and yellow arrows). Not surprisingly, blebbistatin treatment reduced the narrowing of myosin stacks seen in controls (Fig 9H; quantified in Fig 8E). Cells retained the ability to regain a more columnar architecture as lateral borders zipped up, although apical flattening was reduced (Suppl Fig 5A-D).

**Fig 9.**
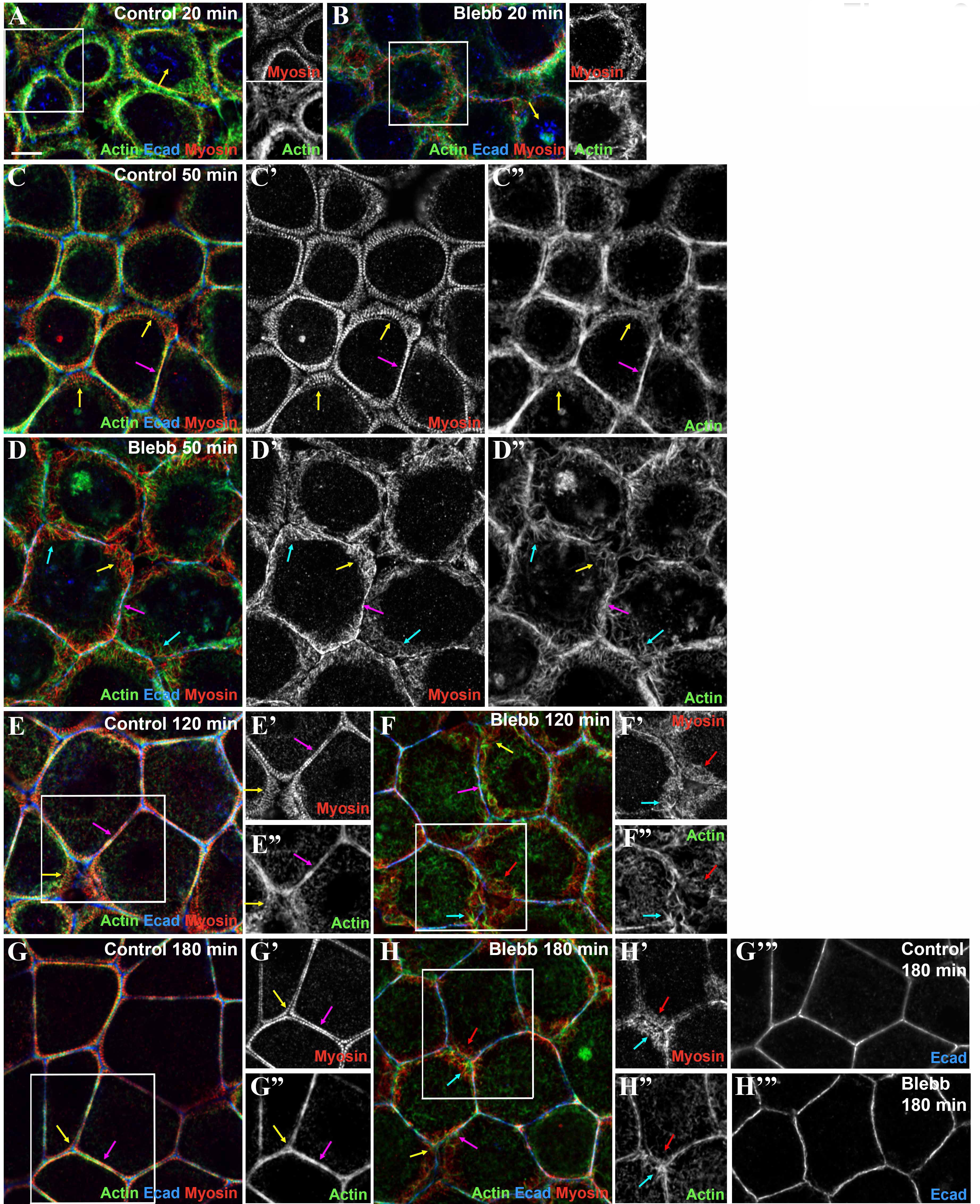
Blebbistatin (Myosin ATPase inhibitor) treatment disrupted assembly of myosin stacks and altered F-actin structures at junctions. (A-G) Representative images, different timepoints during recovery. Control vs. Myosin ATPase -inhibited (Blebb). At midtime points assembly of expansive stacks of myosin is lost (C’ vs D’, arrows), and later in recovery the tight sarcomeric array of myosin at the ZA is lost (E’ vs F’, G’ vs H’). in the presence of Blebbistatin. Actin bundling at the ZA is disrupted and instead actin has a spiky appearance (D”, F”,H”) Also see quantifications in Figure 8E-H.

**Fig 10.**
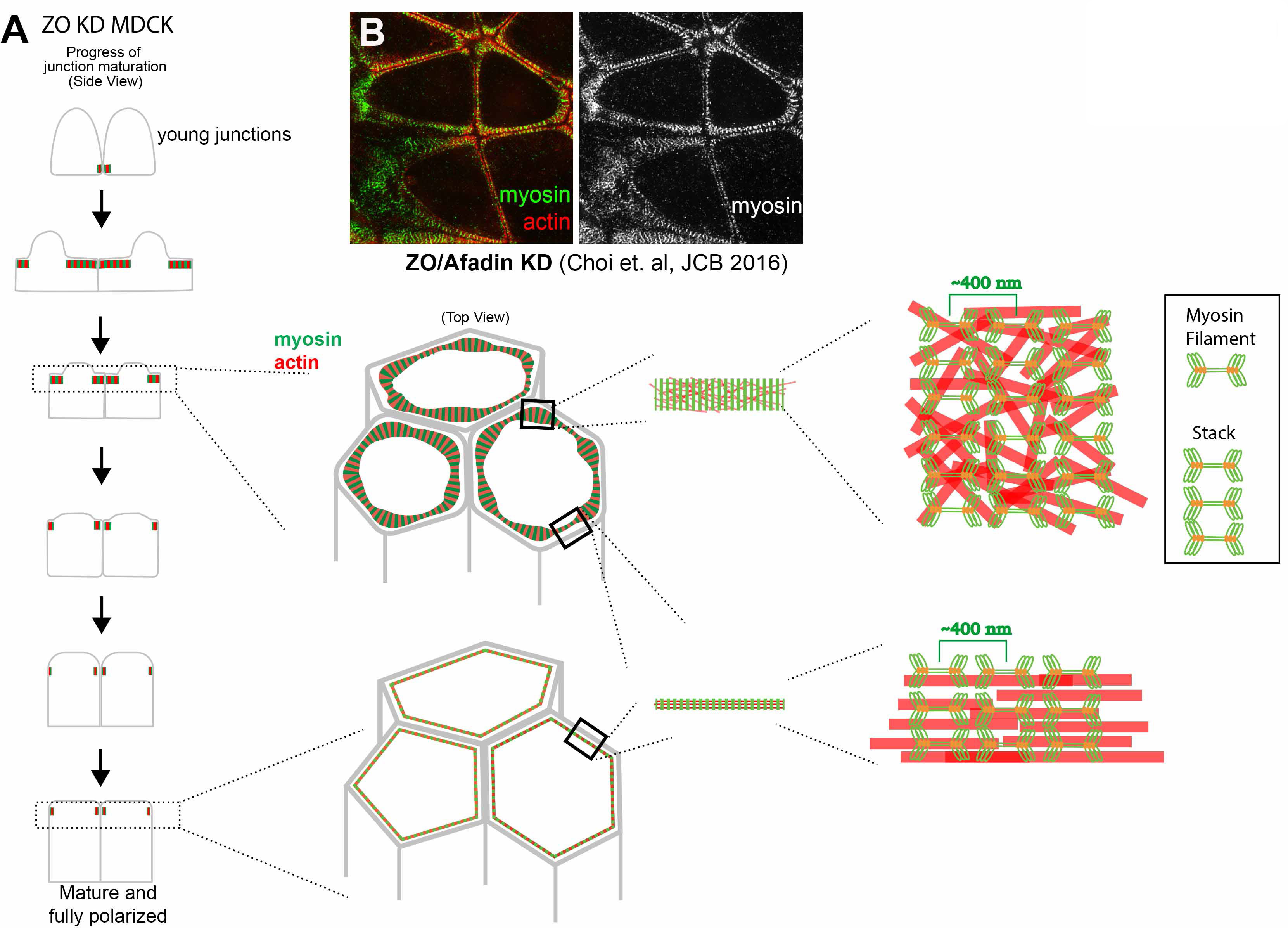
A. Summary Diagram illustrating our model. B. Broad actin stacks assemble when Afadin is knocked down in ZO-knockdown MOCK cells.

After blebbistatin treatment, Ecad, which was endocytosed after calcium removal (Fig 9A, B arrows), still returned to cell borders, where it associated with a junctional population of actin. However, junctional actin architecture was dramatically altered by blebbistatin treatment. This began early in recovery (Fig 9B) and continued to the end of the process (Fig 9H). In controls at the recovery midpoint, extended myosin stacks (Fig 9C’, yellow arrows) overlaid a disordered meshwork of actin filaments (Fig 9C”, yellow arrows), and as myosin stacks narrowed, beginning at bicellular borders (Fig 9E’, magenta arrow), actin filaments became increasingly concentrated at nascent junctions, leading to assembly of bundled arrays (Fig 9E”, G”, arrows). In contrast, in blebbistatin-treated cells spiky actin filaments emanated from cell junctions. These spiky filaments were already evident within 5 min of recovery (Fig 9B, inset), become prominent at mid-stages (Fig 9D”, cyan arrows), and remained present even late in recovery (Fig 9H”, cyan arrow). Quantification of junctional actin confirmed the failure to tightly bundle actin and its replacement by a disordered apical actin array (Fig 8F). Blebbistatin also prevented actin polarization to the ZA (Fig 8G) and cell border straightening was substantially reduced (Fig 9H; quantified in Fig 8H), suggesting that inability to assemble ZA sarcomeric actomyosin may reduce border contractility.

Strikingly, all these effects were rapidly reversed after blebbistatin washout (which began at 90 min of recovery). Within 15 min some bicellular borders straightened and the highly spiky actin at junctions was reduced (Suppl Fig 5G vs H). Within 90 min most borders had assembled sarcomeric myosin and tightly bundled actin (Suppl Fig 5I) and by 150 min cells resembled controls (Suppl Fig 5J), with bundled actin (Fig 8F) and myosin (Fig 8E) at the ZA and straightened cell borders (quantified in Fig 8H). Consistent with a role for myosin in actin bundling, as recovery proceeded after washout there was a correlation between borders where myosin stacks narrowed and those where actin was bundled (Suppl Fig 5H,I magenta arrows), while borders where myosin remained disorganized retained disorganized cortical actin (Suppl Fig 5H,I yellow arrows). Together these data support the hypothesis that myosin motor activity is critical for assembling the ZA supramolecular contractile actomyosin array, and reveal that inhibiting myosin motor activity triggers striking changes in actin architecture at nascent junctions.

## Discussion

The assembly of polarized supramolecular actomyosin structures at the ZA underlies the remarkable ability of cells to change shape and move during embryonic development and tissue homeostasis. Despite decades of work on this issue, key questions remain about the respective contributions of actin and myosin regulators in this process. We used ZO-knockdown MDCK cells, which assemble a textbook ZA, to define underlying mechanisms. Our data reveal that myosin activity plays a key role in driving actin organization at cell-cell contacts, via self-organization of extensive micrometer-scale stacks of myosin filaments underlying the plasma membrane apical to forming cell junctions, which then go through a process of compaction, driving bundling of actin filaments as cell-cell AJs mature (Fig 10A).

### No single actin polymerization machine is essential for assembling supramolecular actomyosin structures at the ZA

Many previous studies explored roles of actin nucleation and continued polymerization in building and maintaining actin structure at AJs. We initially hypothesized that one or more of the machines driving actin polymerization would be essential. Consistent with earlier work (see Introduction), inhibiting Arp2/3 or formins reduced actin levels at cell junctions. However, our data suggest that, at least in this cell type, individual activities of Arp2/3, formins, or Ena/VASP proteins are not essential for assembling supramolecular actomyosin structures at the ZA, with sarcomeric myosin arrays and tightly bundled actin. Previous work is consistent with the idea that this is true in a broader array of cell types, as most previous perturbations reduced junctional actin by 40-60% but did not eliminate it. One speculative possibility is that different actin polymerization machines act in parallel with one another, and one would need to perturb them together to eliminate junctional actin assembly. This will be important to test. It will also be important to test roles of actin bunding proteins, which may work with myosin to organize and bundle actin filaments.

### Myosin activation and motor activity play key roles in ZA assembly

Instead, our data suggest myosin plays a key role in assembling ordered actin to create tight actin bundles at the ZA. This required both myosin activation and myosin motor activity. In our MDCK cells myosin recruitment to cell borders was regulated by phosphorylation of myosin regulatory light chain, which activates and allows myosin to interact with its binding partners. Myosin motor activity was not required for it to localize to cell junctions, but sarcomeric organization required myosin motor function. After blebbistatin treatment, myosin’s ability to cross-link actin filaments may remain intact, suggesting this activity is not sufficient for myosin to drive junctional actin bundling. Myosin’s ability to integrate existing actin filaments into larger actomyosin bundles has been described in studies of stress fibers in lamella (e.g. (Anderson et al., 2008; Machesky and Hall, 1997; Nemethova et al., 2008), where actin polymerization is slow compared to lamellipodia (Glacy, 1983; Wang, 1984; Zicha et al., 2003).

It’s important to note that neither inhibiting myosin activation nor motor activity prevented reassembly of cadherin-based AJs, as cells zipped up along their lateral borders. Ecad was associated with junctional actin—however, this actin was differently organized and apical polarization of Ecad and actin was reduced.

This suggests multiple mechanisms drive actin assembly at AJs, consistent with recent analysis suggesting that actin at the ZA includes both a central branched actin network and adjacent zones of linear bundled actin (Efimova and Svitkina, 2018). The actin associated with interdigitated Ecad complexes seen at midpoints of junctional reassembly may be generated by one of these mechanisms—perhaps it is Ena/VASP driven, as was suggested in keratinocytes. We also were intrigued by the “spiky” nature of junctional actin in blebbistatin-treated cells. Perhaps myosin motor activity or the actin bundling it promotes inhibits other actin assembly pathways at cell junctions.

### A role for self-organized micrometer-scale stacks of myosin filaments

Perhaps our biggest surprise was to see self-assembly of micrometer-scale stacks of myosin filaments with a sarcomeric organization quite early during junctional reassembly (Fig 10A). These arrays were similar to but more extensive than myosin filament stacks in migrating fibroblasts (Beach et al., 2017; Fenix et al., 2016; Hu et al., 2017; Verkhovsky et al., 1995). It was especially intriguing that myosin organized into this large-scale sarcomere-like pattern at locations where Actin remained much less well-organized, suggesting myosin supramolecular organization does not require pre-organized Actin. This echoes seminal observations by the Borisy group, who noted that in fibroblasts “Some myosin spots and ribbons were found in the zone of diffuse actin distribution, suggesting the formation of myosin spots may precede the organization of actin filament bundles.” (Verkhovsky et al., 1995).

The mechanisms mediating myosin stack assembly and lateral interactions remain unclear. Actin and myosin-binding partners might play roles in myosin’s localization and supramolecular organization. Verkhovsky et al. suggested a role for alpha-actinin (Verkhovsky et al., 1995), as it is observed in alternating A-and I-bands of muscle sarcomeres. Consistent with this, α-actinin-4 knockdown disrupted myosin stack assembly on stress fibers in REF52 cells. Knockdown also suggested roles for Cofilin1 and the formin Fmnl3 (Hu et al., 2017). Another possible myosin-binding partner that might regulate assembly is Myosin-18. Myosin-18 can co-assemble with other myosin isoforms to regulate their biophysical properties and localization. It was proposed that Myosin-18 can anchor mixed filaments to the plasma membrane via binding to PDZ-ligand containing proteins (Billington et al., 2015). Heterotypic filaments of Myosin-18 and Myosin-2 exist both in vitro and in stress fibers of Hela, Rat2 fibroblast or U2OS osteosarcoma cells (Billington et al., 2015; Jiu et al., 2019). Myosin-18B promoted formation of myosin-2 stacks and higher-order structure in contractile stress fibers (Jiu et al., 2019). Perhaps Myosin-18 isoforms are present in our system, helping anchor myosin stacks to the membrane and further assist stack formation.

Mammals have three myosin 2 isoforms, Myo2A, Myo2B, Myo2C. Intriguingly, Myo2B localizes to the central branched actin region while Myo2A localizes along the adjacent linear actin bundles (Heuze et al., 2019). Myo2A and Myo2B knockdown also suggest differential roles; e.g. Myo2A and Myo2B have distinct localizations and functions at Ecad-mediated AJs in parental MDCK cells, a cell line that does not form sarcomere-like actomyosin bundles at AJs (Heuze et al., 2019; Ozawa, 2018; Smutny et al., 2010). Myo2A plays a more central role in vivo: myo2A knockout mice have defects in cell adhesion by E6.5 and die by E7.5 (Conti et al., 2004), while myo2B knockout mice die perinatally with defects in the heart and brain (Tullio et al., 1997). Different isoforms also have distinct dynamic properties (Vicente-Manzanares et al., 2009). Myo2A has the highest rate of ATP hydrolysis and thus can move actin more rapidly, while Myo2B has the highest duty ratio, allowing it to exert force on actin longer. Despite their different biophysical properties, the three isoforms have both unique and redundant cellular localizations and functions (Bao et al., 2005; Conti et al., 2004; Sandquist and Means, 2008) (Beach et al., 2014), and can coassemble into heterotypic filaments (Beach et al., 2014). It will be interesting to explore roles of different myosin isoforms throughout junction maturation, both during formation of micrometer-scale stacks of myosin filaments and of the final robust sarcomeric actomyosin structure. We focused on localizing Myo2B, but we also detect Myo2A in both large myosin arrays and in the final ZA. It will be interesting to determine whether heterotypic myosin filaments form in our cell type, and whether the composition and ratios of heterotypic filaments change over time as actin filaments become bundled, supporting the sorting mechanism proposed by Beach et al. (Beach et al., 2014).

One final issue of interest is the role of Ecad and associated linker proteins in directing ZA assembly. In our previous work we explored the role of the junction-actin crosslinker Afadin in ZO-KD MDCK cells. Afadin knockdown had multiple effects in this cell type (Choi et al., 2016). Cell border contractility homeostasis was disrupted, with some borders hyper-constricted and others hyper-extended. Most intriguing with regard to our results here was that effects were most striking at tricellular and short multicellular junctions, the same places where ZA reassembly was slowest. At these “weak points”, we observed expansion of the “sarcomere-like” array seen in ZO-KD MDCK cells into more extensive myosin stacks (Fig 10B). Thus Afadin knockdown cells resembled those observed here during mid-assembly. These data suggest Afadin helps maintain the tight bundling of actin at the ZA, and in its absence extensive myosin arrays assemble at tricellular junctions. It will be interesting to explore mechanisms by which Afadin acts to regulate actomyosin assembly and homeostasis.

## Acknowledgements

We thank Josh Lawrimore for advice on data analysis and Matlab codes, Stephanie Gupton for sharing reagents, Wangsun Choi, Jordan Beach, Richard Cheney, Steve Rogers, Anja Schmidt and other Peifer lab members including for helpful advice and comments, and Tony Perdue of the Biology Imaging Center. This work was supported by NIH R35 GM118096 to M.P.

## Methods

### Cell line, Reagents and Antibodies

MDCK ZO-1/ZO-2-KD (ZO KD, clone 3B3; (Fanning et al., 2012) cells were cultured in complete media (high-glucose DMEM (Corning) supplemented with 10% FBS and 20 mM Hepes (Corning)), and maintained at 37°C with 5% CO2. For imaging, cells were plated on Matrigel (Corning #356231)-coated coverslips. For transfection of GFP-mito-AP4/FP4 constructs (a gift from Dr. Stephanie Gupton, UNC; (Bear et al., 2000), the PolyJet Transfection Reagent (SignaGen Laboratories) was used per manufacturer’s instructions. Calcium switch buffer (CSB) was prepared by adding 2mM Magnesium dichloride into Magnesium-Calcium-Free PBS (Corning). Antibodies and concentrations used for immunostaining are as follows: rabbit anti-non-muscle myosin IIB-targeted at myosin tail (BioiLegend #909902) 1:300, rat anti-E-Cadherin (Santa Cruz, DECMA-1 #sc59778) 1:300, mouse anti-VASP (ECM Biosciences #VM2771) 1:150, mouse anti-ARP3 (Sigma A5979) 1:100. Secondary antibodies were purchased from Thermo Fisher. Actin staining using FITC-Phalloidin or Alexa-647 Phalloidin (Invitrogen) was done according to manufacturer’s instruction.

### Calcium switch and Immunostaining

Calcium switch experiments were performed as previously described (Gumbiner and Simons, 1986), with minor modifications. Cells were grown 48HR after confluency before going through calcium switch. Cells were washed once with CSB, incubated in CSB for 1.5 hours, and then switched back to complete media, which we referred as the recovery period. Samples were collected and immunostained at different recovery timepoints. For drug treatment, inhibitors were added and incubated with cells during the recovery period. Concentration for inhibitors are as followed: 100 µM CK666 (Sigma), 50µM SMIFH2 (Sigma), 100µM Y-26732 (Sigma), 10µM Blebbistatin (Sigma). For Y-26732 or Blebbistatin washout experiments, complete media without drug was used to replace the drug-containing media 90 min into the recovery period.

### Immunostaining

Formaldehyde fixation was used for most experiments, except Fig 3A and Fig 5A, 5B and 5E. For formaldehyde fixation, 1% of formaldehyde was used to fix cells. Cells were then quenched and permeabilized with 0.5% TritonX-100, then incubated with primary and secondary antibodies. For Fig 3A and Fig 5B, 5E, cells were permeabilized before fixation, using a method described in (Yu-Kemp et al., 2017). For Fig 5A, cells were fixed with ice-cold methanol for 5 minutes, then permeabilized with 0.1% TritonX-100.

### Imaging and Image Processing

Images were collected via Airyscan or SIM imaging. Airyscan imaging was performed on a Zeiss LSM880 system, with a X63 1.4NA oil objective. Z collection with intervals at 0.18 µm. Raw images were processed using the 3D Airyscan processing function in Zen software (Carl Zeiss). SIM images were collected using a Nikon N-SIM and SR APO TIRF with a 100 ×/1.49 NA oil objective with 3D-SIM function. Images were processed with stack reconstruction using NIS-Elements AR Version 4.51 software 2016 (Nikon). Imaging parameters across conditions were kept identical for each set of experiment, to allow comparison of intensity between control and the drug treated conditions. All x-y (top view) images in the figures are maximum intensity projections of the apical region of the cell, unless otherwise noted. 3D images were generated using Imaris 9.6 (Bitplane) with the surface function. To visualize myosin and cadherin signals at the border in the surface view, the actin signal at the border at the borders was selected with surface gain size of 0.3-0.5 µm and then deducted from the total actin surface area. Photoshop CS6 (Adobe) was used to adjust input levels so that the signal spanned the entire output grayscale and to adjust brightness and contrast.

### Quantification

Quantifications were done using Fiji (NIH) or MATLAB. Individual cells or borders for measuring signal intensity or border curvature were manually selected based on the 3x3 grid in Fiji/ImageJ (National Institutes of Health). Our approaches are illustrated diagrammatically in Suppl Fig 6. For measuring myosin covered area, the myosin signal was binarized, and value of pixels that are covered by myosin were extracted using Matlab. For measuring myosin sarcomere spacing, a straight line was drawn at the cell-cell border in Fiji, the intensity value was extracted and the mean interval between maxima, which indicates the sarcomere spacing, was calculated by MATLAB using the findpeaks function. For measuring border curvature, the length of a line drawn from fully tracing the cell border from one tricellular junction to the next and compared to the length of a straight line drawn directly from one tricellular junction to another. Only borders with a straight-line length between 150 and 450 pixels (6.4 to 19.2 µm) were measured. For measuring actin and cadherin intensity, borders were surrounded by an 8- x 3.5-µm box and the intensity values were measured at the apical z-slice and the lateral z-slice, with lateral defined as halfway between the apical z-slice and basal z-slice. For measuring actin covered area borders were surrounded by a 7.5- x 3-µm box. Each box was binarized and the value of the pixels corresponding to actin covered area were extracted using the Histogram tool. All statistical analyses were performed using Prism 8 (GraphPad Software). Fitted curves for myosin area ratio or border straightness were calculated using non-linear fit in Prism. Scatter plots presents all the quantified data points, with mean ± SD.

---

**Suppl Fig 1.**
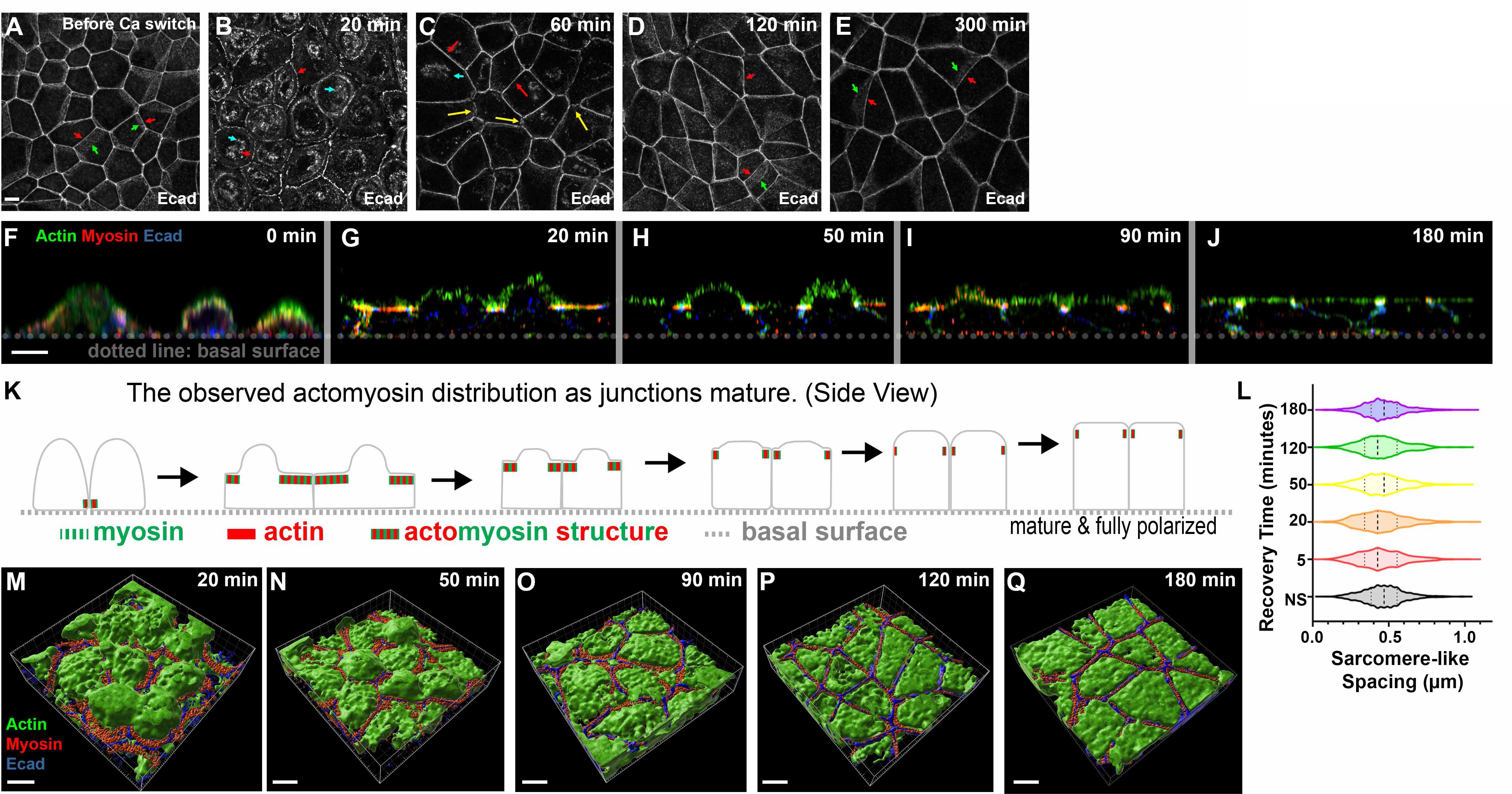
(A-E) The localization of Cadherin throughout the process of calcium recovery. Cadherin is internalized right after the calcium switch (cyan arrows). As the recovery proceeds, Ecad regained its continuous distribution at the ZA, and cell borders straightened. Arrows: (red) apical ZA; (green) lateral membrane; (yellow) tricellular junctions. (F-J) Cross-section view of monolayer as junctions mature. The apical membrane flattened as cells polarized. Myosin is enriched in arrays underlying the apical membrane which narrow until it is enriched at the ZA. (K) Diagram illustrating actomyosin localization along the Z-axis as junctions mature. (L) Quantification of the myosin sarcomere-like spacing at different recovery timepoints. The spacing remained ∼0.45 µm throughout junction maturation, indicating the age of the junction did not modulate myosin spacing. NS=Non-switched. (M-Q) Three-dimensional surface view of cells at different recovery timepoint. These images reveal how the apical surface of the cell changed as junctions and myosin structure matured.

**Suppl Fig 2.**
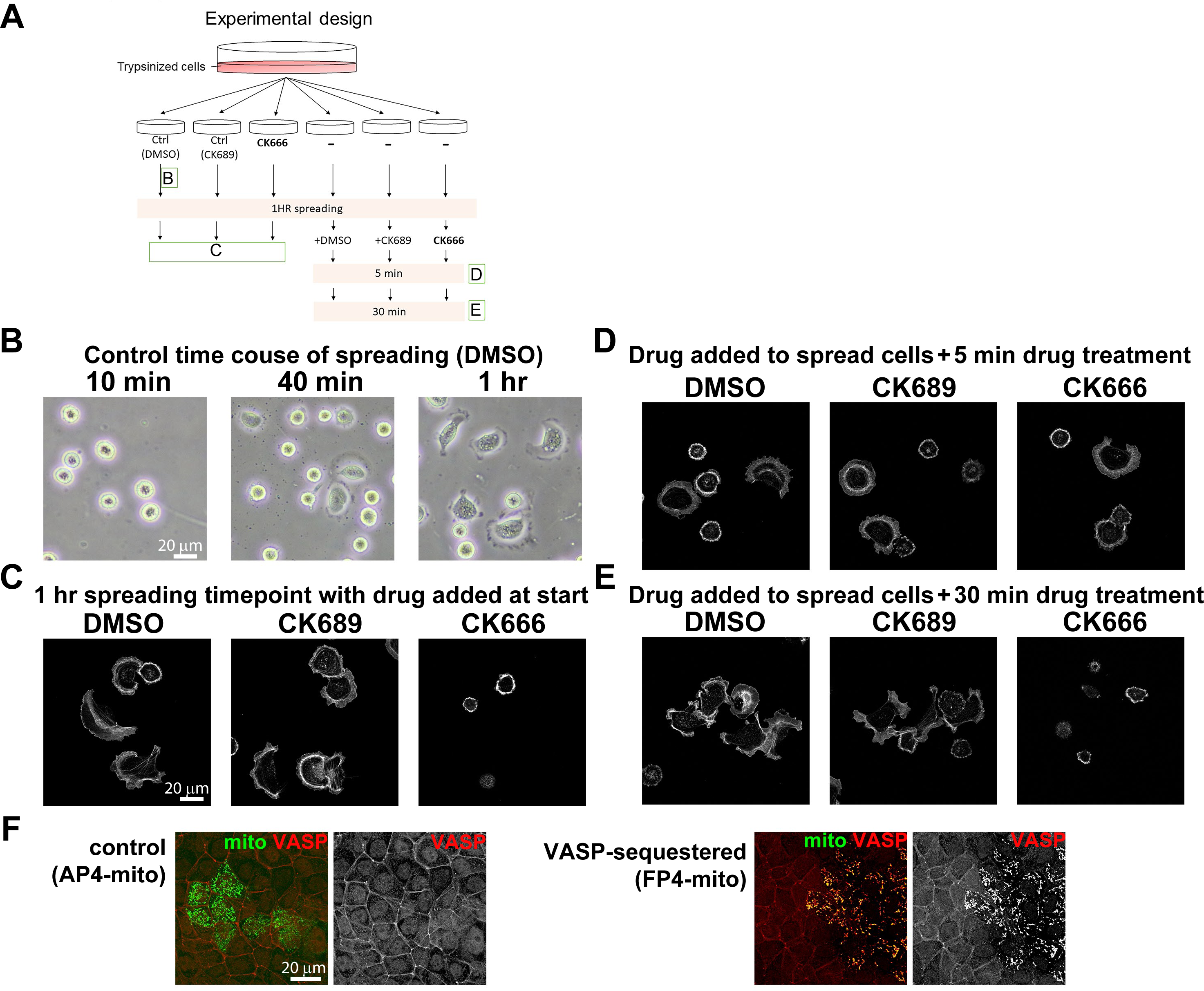
Verification of the activity of CK666 for inhibiting Arp2/3 function and of AP4/FP4 constructs for sequestering Ena/VASP to mitochondria. (A-E) Verification of CK666 function. (A) Schematic diagram of the experiment. (B) Bright field image of control cells at different spreading timepoints. More Arp2/3-dependent lamellipodia structures were formed the longer the cells were plated on extracellular matrix. (C-E) Phalloidin staining. (C) CK666 inhibited Arp2/3-dependent cell spreading, whereas the control (DMSO or CK689 (the inactive molecule) did not. (D-E) Arp2/3-dependent lamellipodia disappeared in the presence of CK666. (F) Verification of VASP-sequestering construct. In cells transfected with the control construct (AP4-mito) VASP remained enriched at the cortex (left panel). In FP4-mito transfected cells VASP relocalized to internal structures we presume are mitochondria (right panel).

**Suppl Fig 3.**
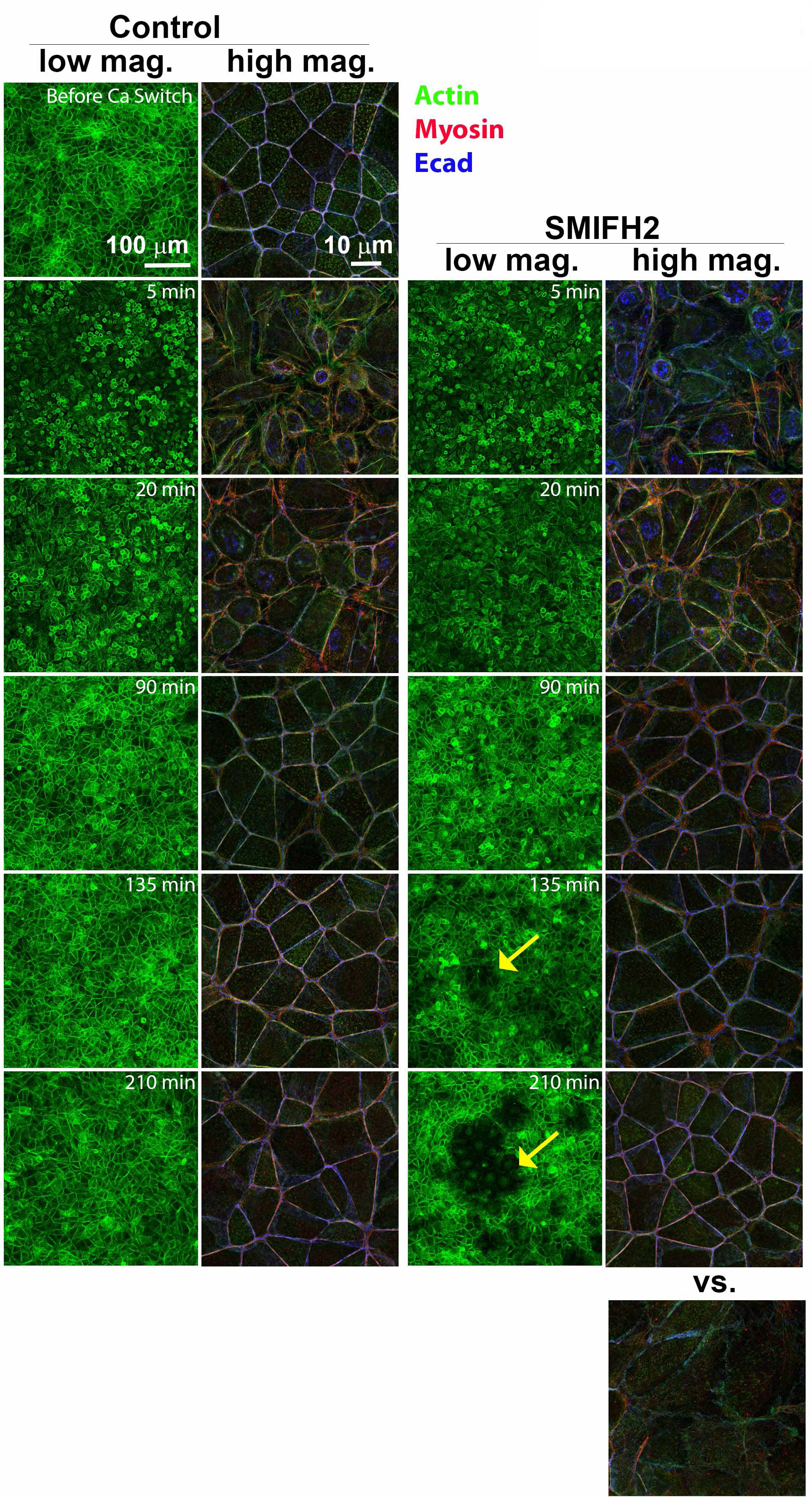
Images demonstrating the effect of SMIFH2 on a ZO KD MOCK monolayer. When the cells recovered from calcium switch **in** the presence of SMIFH2, we observed regions **in** the monolayer with reduced actin signal (yellow arrows) and abnormal cell shape. The longer the cells were incubated with SMIFH2, the more noticeable these “defects” were **in** a monolayer.

**Suppl Fig 4.**
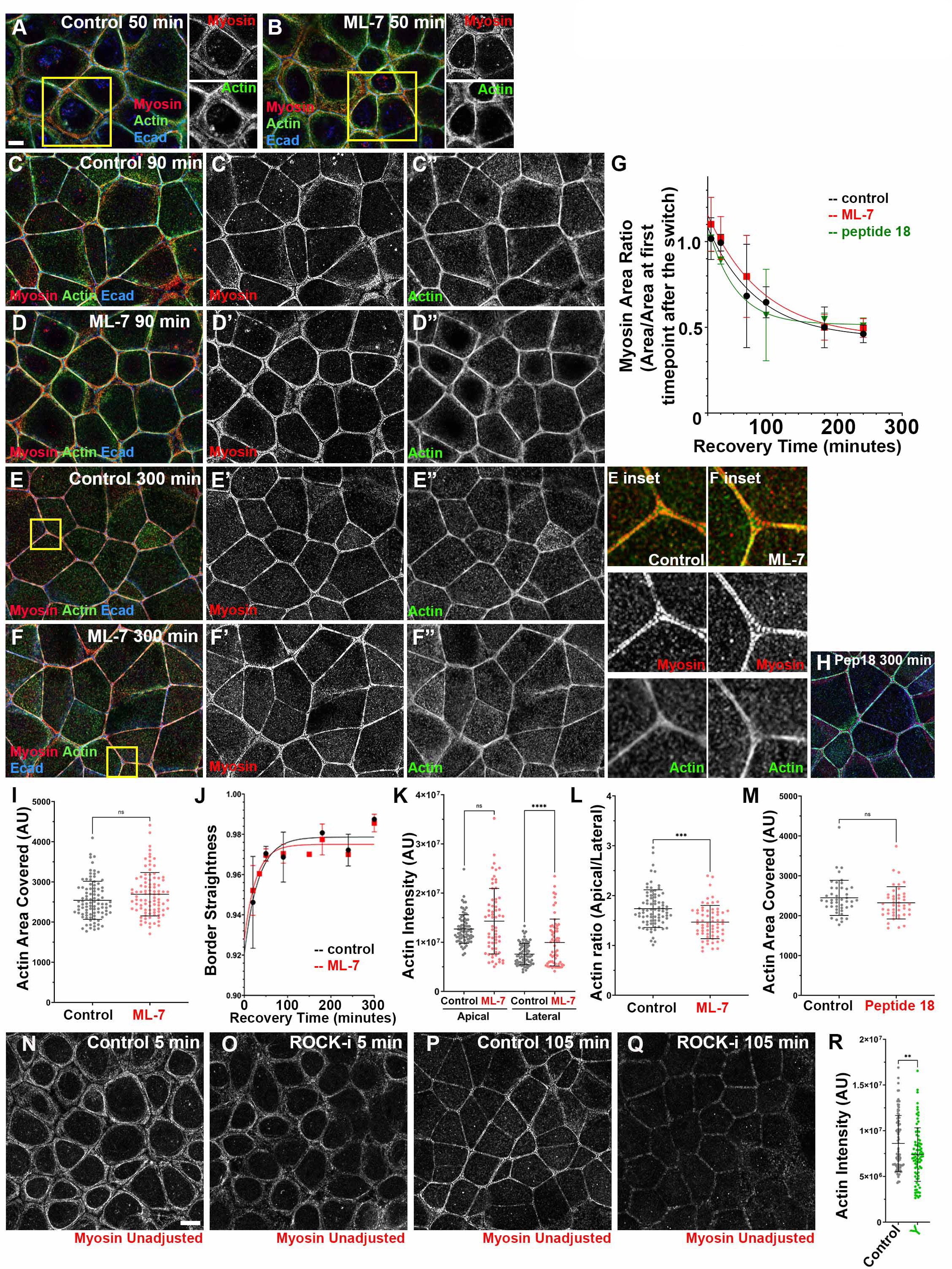
Inhibiting MLCK using ML-7 or peptide-18 does not alter myosin localization during recovery or prevent or delay assembly of the final ZA actomyosin structure. (A-F) Representative images showing cells in control vs. ML-7-treated cells at different timepoints during calcium recovery. By the last timepoint, cells assemble bundled F-actin decorated by sarcomeric myosin in both control and drug-treated conditions (E vs F, H). (G, I-M) Quantification. Myosin maturation (G), actin bundling at the ZA (I,M), and border straightening (J) were similar in control and drug-treated conditions. After ML-7 treatment here was some elevation of lateral myosin and thus apical actin polarization was reduced (L) . (N-Q) Representative images showing the difference in apical myosin signal in control (N, P) vs ROCK-inhibited (O, Q) conditions. Myosin signal at the junction reduced when ROCK is inhibited. (R) Quantification. Levels of apical actin were reduced after ROCK-inhibition.

**Suppl Fig 5.**
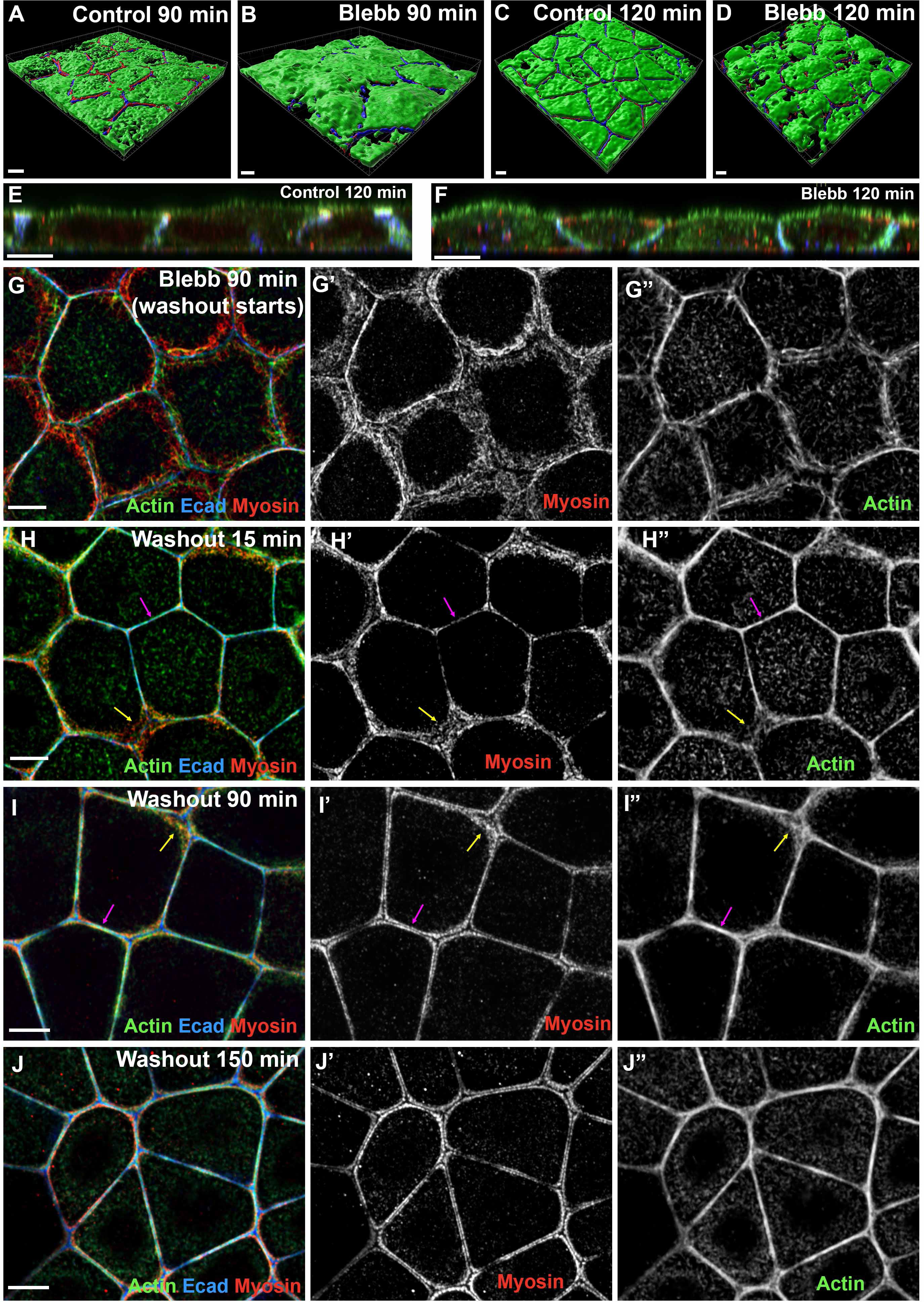
(A-D) Three-dimensional images of cell surface in control vs. Blebbistatin-treated cells. The apical surface of the control cells becomes flatter as junctions mature (A vs C, also in Suppl Fig 1M-Q). In contrast, Blebbistatin-treated cells were not able to flatten the apical surface, even at the later recovery timepoints (B and D). (E-F) Cross-section views. After blebbistatin treatment, cells still retained the ability to regain a more columnar architecture as lateral borders zipped up, even though actin, myosin and Cad were not apically polarized (E vs F). G-J. Representative images showing the recovery of ZA actomyosin structures after Blebbistatin washout. Within 15 minutes after washout, myosin already began to focus at at bicellular borders (H’ and H’’, magenta arrows). Myosin at the tricellular borders was slower to recover (H, I, yellow arrows) but by 150 min (J), ZA actomyosin structures returned to that seen in the control mature monolayer.

**Suppl Fig 6.**
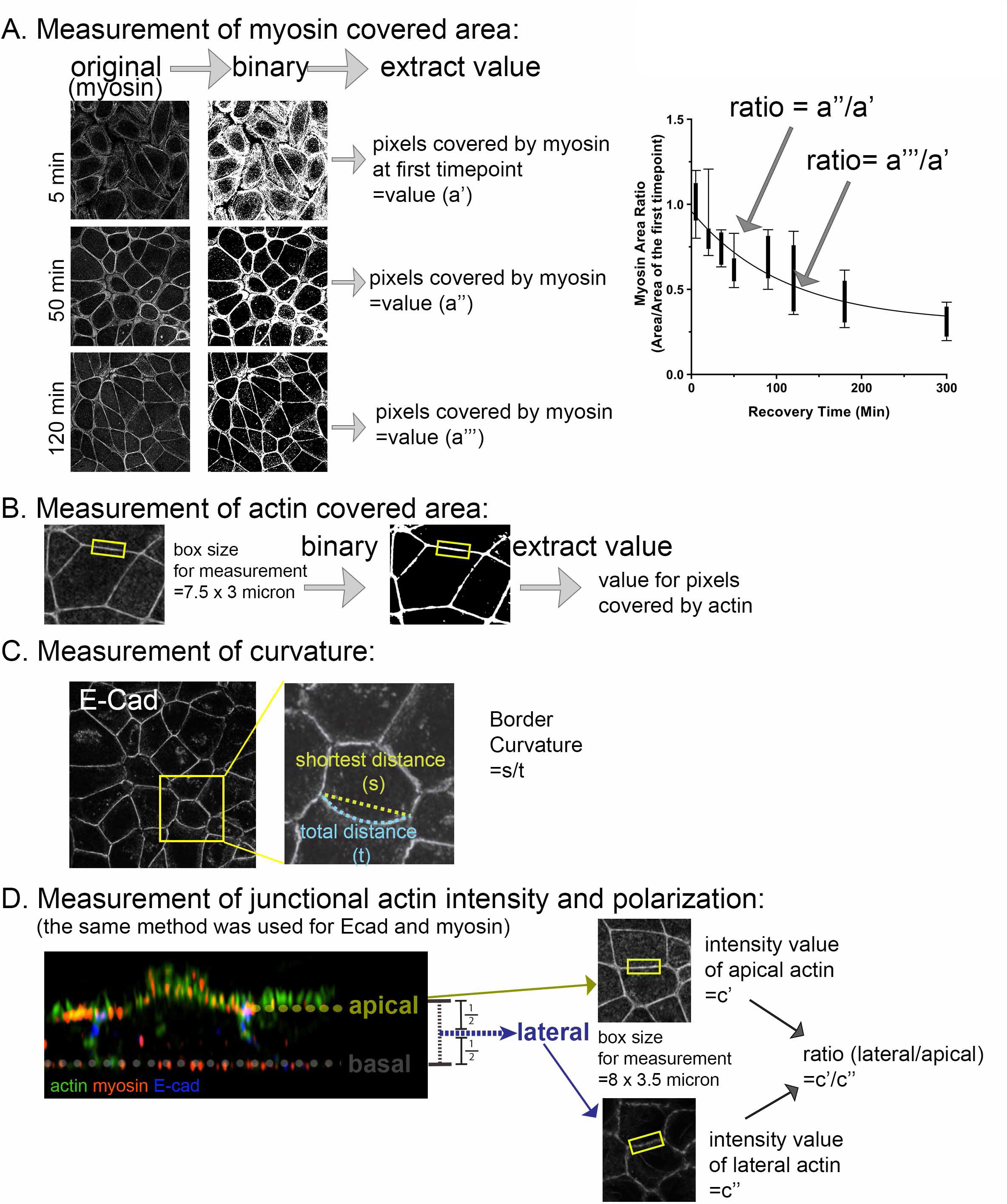
Visual illustration of quantification methods used. Details are in the quantification section of the Methods.

